# The primordial knot: the deep-rooted origin of the disulfide-rich spider venom toxins

**DOI:** 10.1101/2022.10.06.511106

**Authors:** Naeem Yusuf Shaikh, Kartik Sunagar

**Affiliations:** Evolutionary Venomics Lab, Centre for Ecological Sciences, Indian Institute of Science, Bangalore 560012, Karnataka, India

**Keywords:** Spider venom, disulphide-rich peptides, venom evolution, toxin superfamily

## Abstract

Spider venoms are a complex concoction of enzymes, polyamines, inorganic salts and disulfide-rich peptides (DRPs). Although DRPs are widely distributed and abundant, their evolutionary origin has remained elusive. This knowledge gap stems from the extensive molecular divergence of DRPs and a lack of sequence and structural data from diverse lineages. By evaluating DRPs under a comprehensive phylogenetic, structural and evolutionary framework, we have not only identified over 70 novel spider toxin superfamilies but also provide the first evidence for their common origin. We trace the origin of these toxin superfamilies to a primordial knot - the ‘Adi Shakti’ - nearly ∼375 MYA in the common ancestor of Araneomorphae and Mygalomorphae. As these lineages constitute over 50% of the extant spiders, our findings provide fascinating insights into the early evolution and diversification of the spider venom arsenal. Reliance on a single molecular toxin scaffold by nearly all spiders is in complete contrast to most other venomous animals that have recruited into their venoms diverse toxins with independent origins. Moreover, by comparatively evaluating araneomorph and mygalomorph spiders that differentially depend on their ability to secrete silk for prey capture, we highlight the prominent role of predatory strategies in driving the evolution of spider venom.

**Significance Statement:** Venoms are concoctions of biochemicals that function in concert to incapacitate prey or predators of venom-producing animals. Most venomous animals secrete a complex venom cocktail, constituted by toxins with independent evolutionary origins. In complete contrast, we trace the origin of diverse toxin superfamilies in spiders to a single molecular scaffold. The common origin of these disulphide-rich peptides that constitute three-quarters of nearly all spider venoms, therefore, represents a unique scenario of weaponization, where a single motif was recruited and extensively diversified to generate a plethora of superfamilies with distinct activities. Remarkably, the evolution of spider venom was also found to be driven by prey capture (i.e., reliance on silk versus venom) and venom deployment (predation or self-defence) strategies.

## Introduction

With their killer instinct and deadly toxins, spiders have been at the centre of many myths and folktales from times immemorial. They are an archetypal arthropod group with mid-Cambrian or early Ordovician origin, nearly 495 million years ago (MYA) (1). Because of their unique ability to secrete silk and venom, spiders have successfully colonised diverse ecological niches. They are amongst the most successful predators on the planet, with over 50,000 species and 129 families described to date (2, 3). The majority of spiders are equipped with chelicerae harbouring venom glands, with Symphytognathidae, Uloboridae, and certain primitive Mesothelae species being the only exceptions (2, 4).

Spider venoms are a concoction of enzymes, polyamines, nucleic acids, inorganic salts and disulfide-rich peptides (DRPs) (5, 6). They are predominantly rich in DRPs that are characterised by a diversity of structural motifs, including Kunitz (7), disulfide-directed β-hairpin (8), disulfide-stabilised antiparallel β-hairpin stack (DABS) (9) and inhibitor cystine knot (ICK) (10, 11). Despite the fact that DRPs constitute three-quarters of the spider venom, our evolutionary understanding of their origin and diversification has remained elusive. This knowledge gap stems from a lack of sequence and structural data for DRPs from diverse spider lineages and the prevalence of significant sequence divergence in these toxins.

Here, we examined DRP sequences from the Mygalomorphae infraorder and the Retrolateral Tibial Apophysis (RTA) Araneomorphae, which constitute over 52% of spider genera (2,200 genera) described to date. A molecular phylogenetic framework implemented in this study resulted in the identification of over 70 novel toxin superfamilies and suggests a deep-rooted origin of venom DRPs in spiders. Our findings also highlight the role of distinct prey capture strategies of Araneomorphae and Mygalomorphae in shaping the recruitment and diversification of venom DRPs. Furthermore, by comparatively evaluating spider venom toxins employed for anti-predatory defensive and prey capture, we also unravel the impact of the purpose of venom deployment on the evolution of spider venoms. Thus, sequence, phylogenetic, structural and evolutionary assessments in this study have provided insights into the fascinating origin and early diversification of this predominant spider venom component.

## Results

### Novel spider toxin superfamilies

Superfamilies (SF) of venom toxins in spiders have been classified based on their signal peptide and propeptide sequences (12). This premise was first used to describe the Shiva superfamily of toxins from Atracidae spiders (12). Recently, using a similar approach, 33 novel spider toxin superfamilies have been identified from the venom of the Australian funnel-web spider, *Hadronyche infensa* (9). Since gene phylogenies have not been extensively utilised while classifying spider venom toxins, our understanding of their origin and diversification has been severely limited.

In this study, we relied on the strong conservation of signal peptide and propeptide sequences in identifying several novel spider venom toxin superfamilies. Blast searches were used to identify the homology between largely divergent toxin superfamilies. Toxin sequences were found to share strong sequence conservation within a superfamily. Cysteine residues, which are involved in the formation of disulphide bonds and, thereby, are extremely vital in determining protein structure and function, were used as guides to manually refine sequence alignments. This approach enabled the identification of 33 novel toxin superfamilies along the breadth of Mygalomorphae (Figures S1 and S2). Among these, 31 superfamilies belonged to the DRP class, whereas, the other two were enzymatic non-DRP toxins, including the first report of Neprilysin (SF103) and CAP (CRiSP/Allergen/PR-1; SF104) from Atracidae spiders (Dataset S1).

Moreover, analyses of Araneomorphae toxin sequences using the strategy above resulted in the identification of 38 novel toxin superfamilies from Araneomorphae, all of which belonged to the DRP class of toxins (Figures S3 and S4). Overall, among all novel spider toxin superfamilies identified in this study, the majority (n=69) were DRPs, reinstating the dominance of this toxin type in spider venoms. Based on the arrangement of cysteine residues involved in the formation of disulphide bonds, these DRPs could be further segregated into ICK-like (n=26), DABS (n=13) and novel disulphide patterned non-ICK (n=30) superfamilies (9).

The identification of novel toxin superfamilies was further supported by phylogenetic and principal component analyses. Reconstruction of evolutionary histories using Bayesian inference (BI) and maximum-likelihood (ML) approaches retrieved monophyletic clades of toxin superfamilies (Figures 1 and 2; node support: ML: >80/100; BI: >0.95; refer to figures S5, S6 and S9 for complete phylogeny with branch lengths). Interestingly, the plesiotypic DRP scaffold seems to have undergone lineage-specific diversification in Mygalomorphae, where the selective diversification of the scaffold has led to the origination of novel toxin superfamilies corresponding to each genus (Figure 1). In our Bayesian and maximum-likelihood phylogenetic tree reconstructions, these toxin scaffolds were found to form distinct monophyletic clades, further supporting this claim (Figure S5; node support: ML: ML: >80/100;; BI: >0.95).

**Figure 1.**
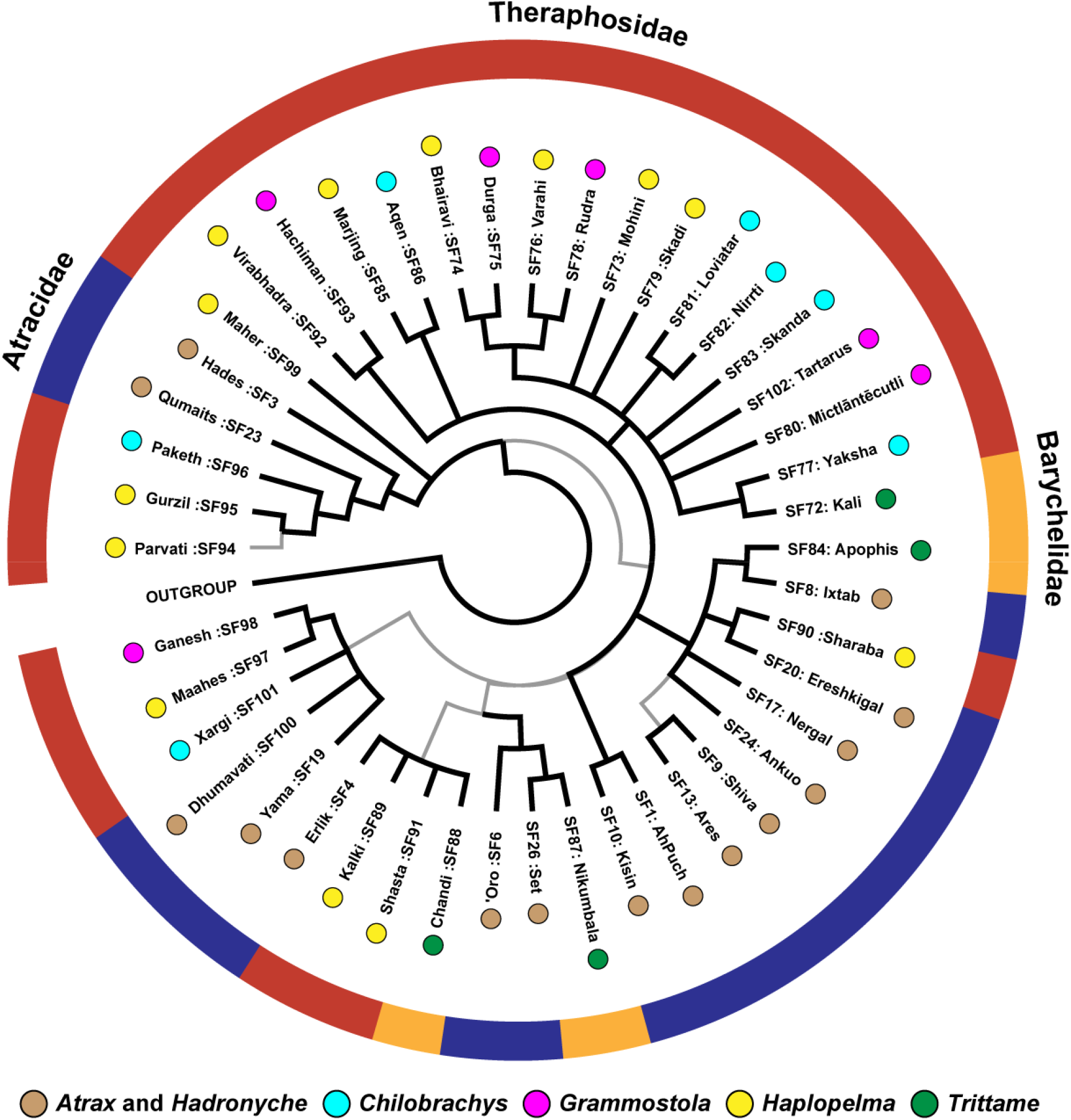
The Bayesian phylogeny of mygalomorph spider venom toxin superfamilies This figure represents the Bayesian phylogeny of Mygalomoprhae spider toxin superfamilies. Coloured spheres alongside tree tips represent the spider genera, while the coloured outer circle indicates the spider family in which the respective toxin superfamily has been identified [Atracidae (red), Barychelidae (orange) and Theraphosidae (blue)].

**Figure 2.**
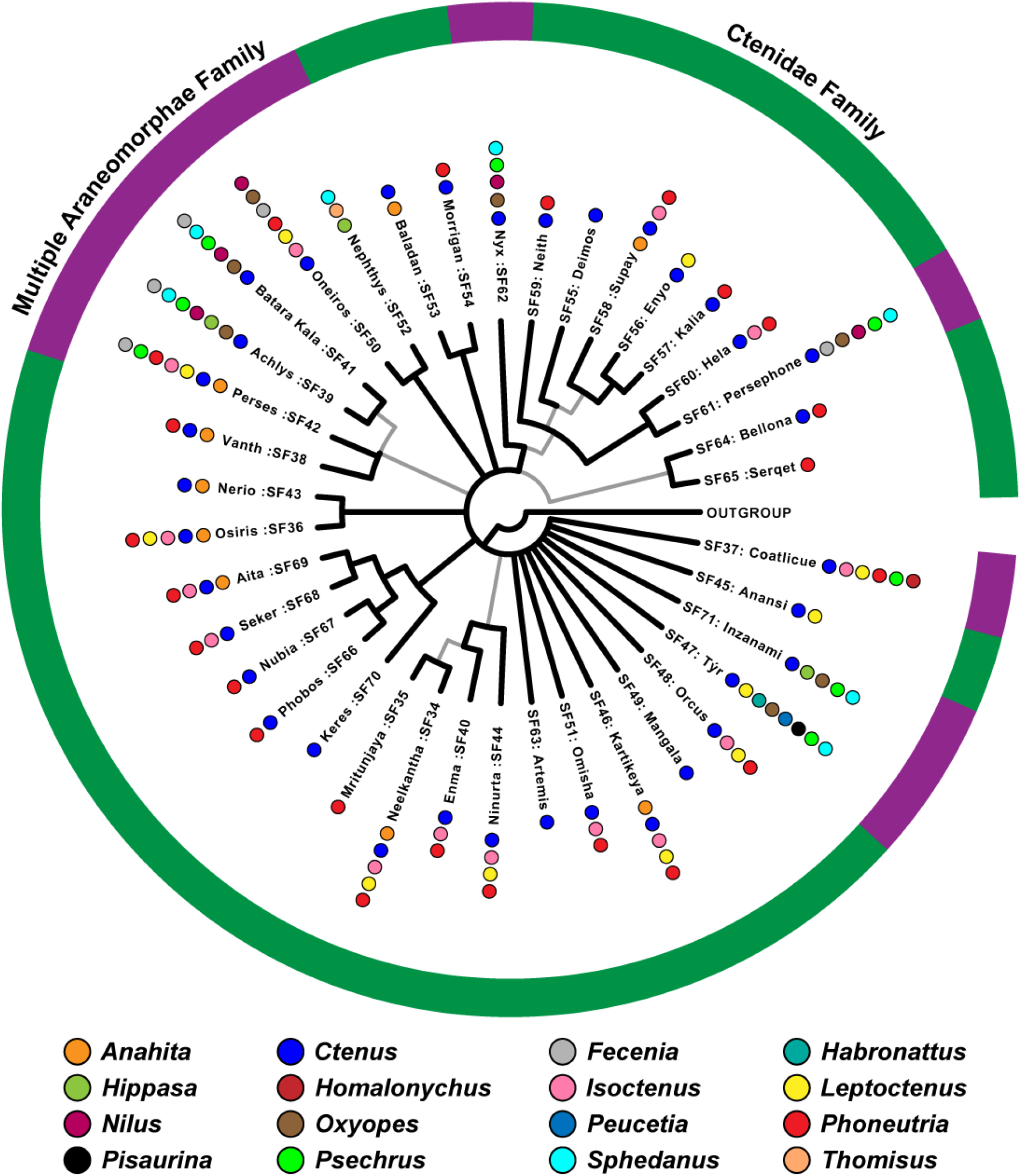
The Bayesian phylogeny of araneomorph spider venom toxin superfamilies This figure represents the Bayesian phylogeny of Araneomorphae spider toxin superfamilies. Coloured spheres, alongside tree tips, represent the spider genera, while the coloured outer circle indicates the spider family [Ctenidae (green), multiple araneomorph families (purple)] in which the respective toxin superfamily has been identified.

A similar pattern was also observed in the case of Araneomorphae, where certain toxin SFs (n=6) were found to have diversified within individual genera (Figure 2). However, we also documented a large number of DRP toxins (n=32) that were found to have diversified in a family-specific manner, wherein, a toxin scaffold seems to be recruited at the level of the spider family, rather than the genus. As a result, and in contrast to mygalomorph DRPs, araneomorph toxin superfamilies were found to be scattered across spider lineages (Figure 2; Figure S6; node support: ML: >80/100; BI: >0.95). Moreover, Principal component analysis (PCA) of toxin sequences further provided evidence for the monophyly of mygalomorph and araneomorph SFs, where each toxin superfamily formed a distinct group in PCA plots (Figures S7 and S8).

Furthermore, sequence alignments of DRPs clearly highlighted the homology among DRP toxin superfamilies (Figure 3; Figure S9; node support: ML: >50/100; BI: >0.95). Six cysteine residues were found to be nearly universally conserved across 101 DRP toxin SFs (Figure 3b; Figure S10). Our findings enabled us to trace the origin of spider venom DRPs in Opisthothelae, the most recent common ancestor (MRCA) of Araneomorphae and Mygalomorphae (13). Thus, we highlight for the first time that all DRP toxins in spiders may have had a common molecular origin, nearly 375 MYA. It should be noted, however, that functional analyses have been performed only on a handful of mygalomorph toxins, with even fewer studies focusing on araneomorph toxin superfamilies, and that it would be inaccurate to speculate the functions of these toxins based on homology.

**Figure 3.**
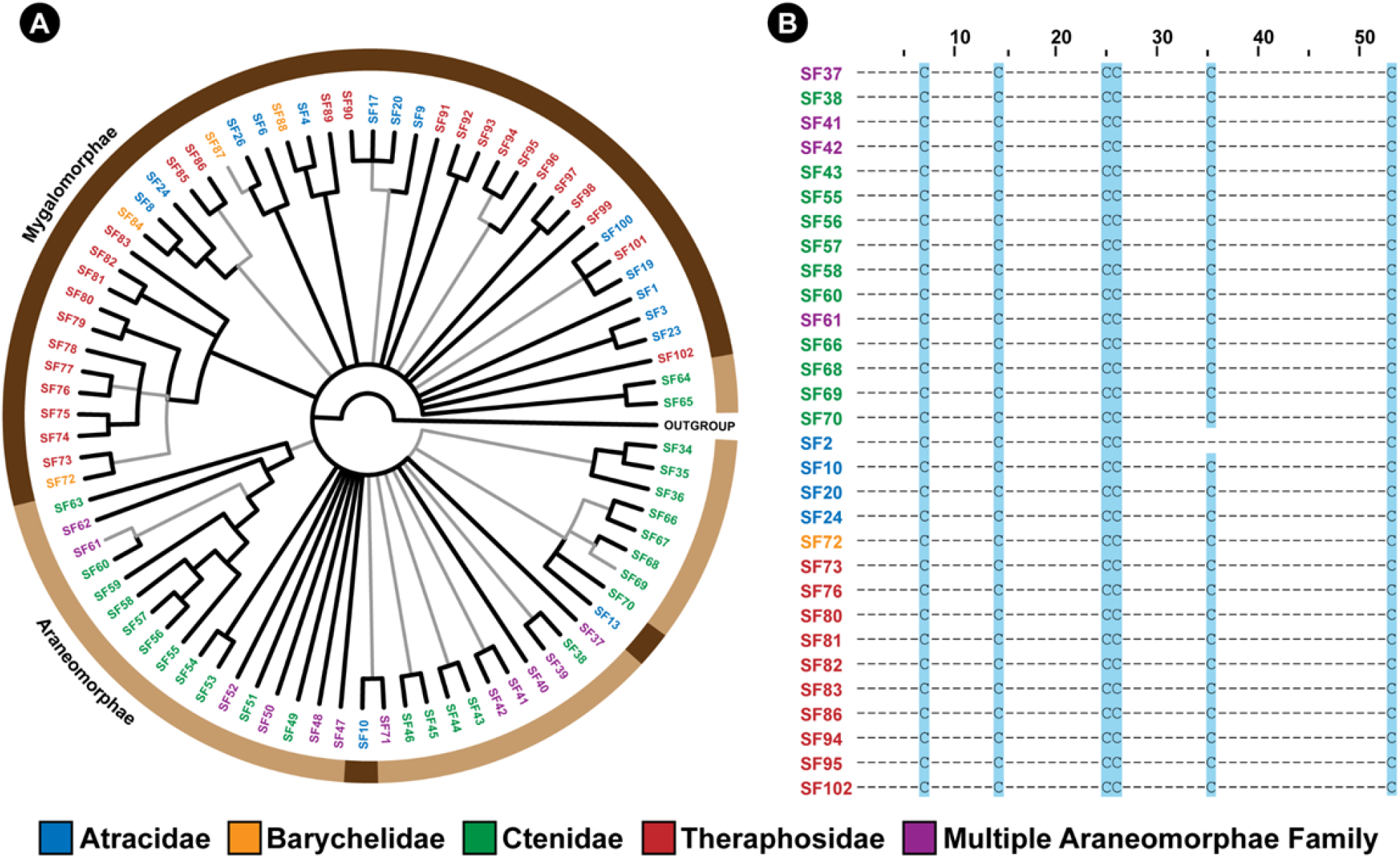
The Bayesian phylogeny and cysteine sequence alignment of spider venom DRPs This figure depicts the Bayesian phylogeny and alignment of representative sequences of Araneae DRP toxin superfamilies. The coloured outer circle in panel A indicates the infraorder of spiders (Mygalomorphae and Araneomorphae shown in dark and light brown, respectively) in which the respective DRP superfamily was identified. In panel B, cysteine residues that are conserved across toxin SFs are highlighted in blue.

### Molecular evolution of spider venom DRP toxins

To evaluate the nature and strength of the selection that has shaped spider venom DRPs, we employed site-specific models that detect selection across nucleotide sites. Our findings suggest that the majority of Mygalomorphae toxin superfamilies (12/19 SFs) have evolved under the influence of positive selection [ω ranging between 1.1 to 2.9; positively selected sites (PS): 0 to 26], while the remaining few have experienced negative or purifying selection (ω ranging between 0.7 to 0.8; PS: 0 to 13; Figure 4, Table S1). In stark contrast, nearly all of the Araneomorph toxin superfamilies that we investigated here were found to have evolved under a strong influence of negative selection (ω ranging between 0.2 to 1.0; PS: 0 to 10; Figure 4, Table S2). We further assessed whether these changes documented across sites have a significant effect on the biochemical and structural properties of amino acids using TreeSAAP (Tables S1 and S2). Outcomes of these analyses revealed the accumulation of replacement changes in Mygalomorphae toxin superfamilies that result in radical shifts in amino acid properties, potentially influencing their structure and function (Table S1).

**Figure 4.**
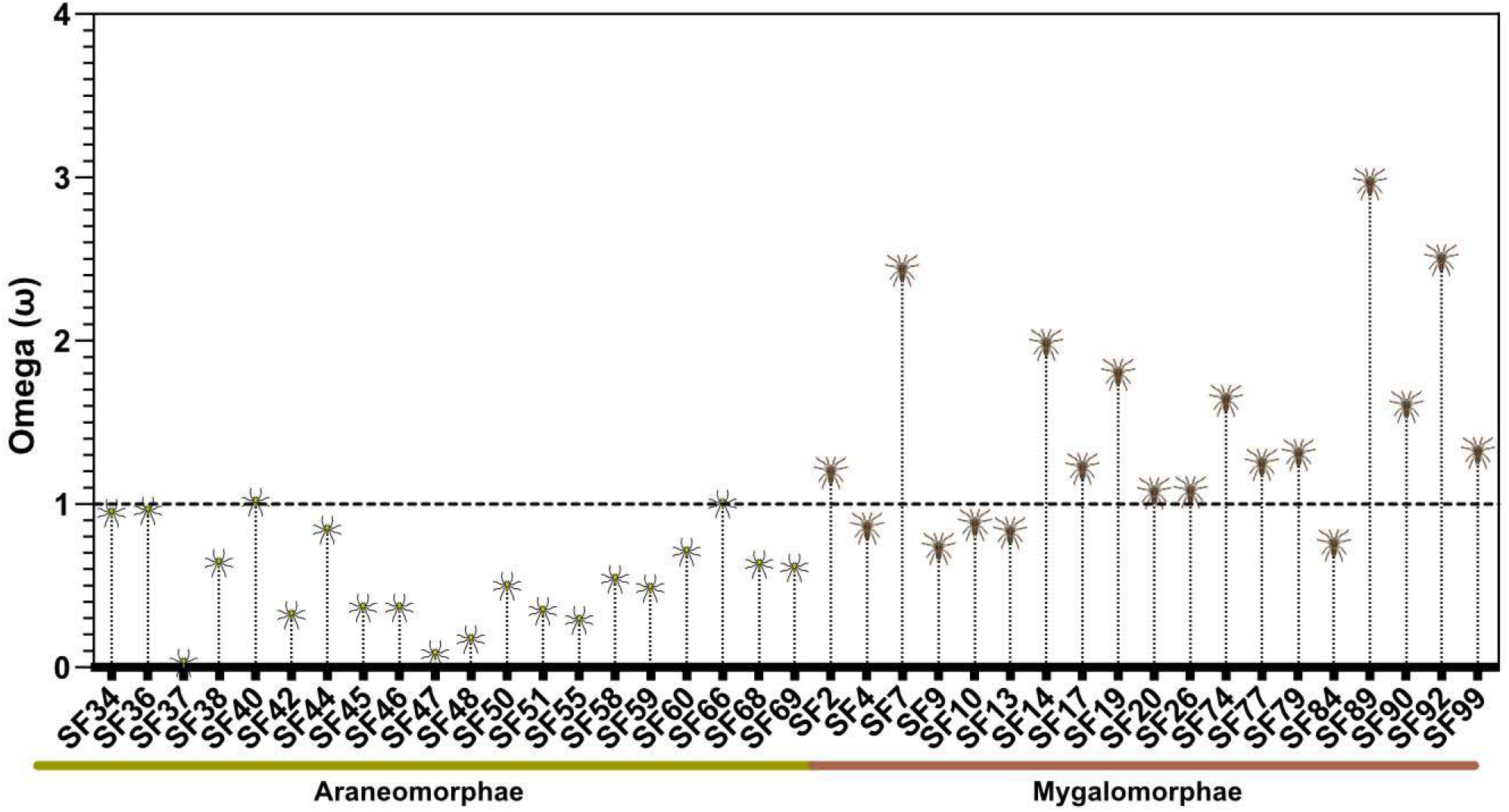
Molecular evolution of spider toxin superfamilies This figure shows the distribution of ω values (Y-axis) for araneomorph and mygalomorph spider venom toxin superfamilies (X-axis). The horizontal dotted red line represents neutral evolution (ω=1), with ω values above and below it indicating positive (ω>1) and negative (ω<1) selection, respectively.

To comparatively evaluate the nature of selection that shapes venom components deployed either for prey capture or antipredator defence, we employed maximum-likelihood and Bayesian approaches. In these analyses, we identified toxin superfamilies SF74, SF77, SF79, SF89, SF90, SF92 and SF99 as predatory toxins (i.e., toxins deployed for prey capture - refer to the discussion section for the principle considered for this classification), whereas SF13 (i.e., Ares SF) was classified as a defensive spider venom toxin superfamily (i.e., toxins deployed for antipredator defence) as described previously (14). Assessment of molecular evolutionary regimes identified a significant influence of positive selection on venom toxins that are employed for prey capture (ω ranging between 1.2 to 2.9; PS: 0 to 11, Table S1, Figure S11), relative to those that are chiefly or exclusively used for antipredatory defence (ω=0.8; PS: 3; Table S1, Figure S11).

## Discussion

### The deep evolutionary origin and diversification of the primordial knot

Prior attempts to explore the phylogenetic and evolutionary histories of spider venom DRPs have hypothesised independent origin and lineage-specific diversification of DRP venom toxins (15). In contrast, recent literature, primarily focusing on *Hadronyche infensa*, suggests that the diverse disulfide-rich venom arsenal of this Australian funnel-web spider is a derivative of an ancestral ICK motif that underwent several rounds of duplication and diversification (9). Often restricted to a specific spider lineage, or given the inconsistent ways of classifying spider venom toxins, previous attempts have failed to provide a broader perspective on the evolution of these peptides (16, 17). Given their very long evolutionary histories, genes encoding DRP toxins have undergone significant diversification, making it difficult to precisely trace their phylogenies. Together with the lack of structural and functional data for these toxins, all of the aforementioned factors have impeded our understanding of the origin and evolution of this predominant spider venom component.

To address this knowledge gap, we employed sequence comparisons, phylogenetic inferences and evolutionary analyses. Our findings strongly suggest a deep-rooted origin of DRP spider venom superfamilies, possibly from a single ancestral DRP or knottin scaffold, which we name ‘Adi Shakti’, the original creator of the universe according to Hindu mythology (Figure 3a). We propose that all of the extant spider toxin superfamilies in Mygalomorphae and Araneomorphae (n=102), which include those that were previously reported (n=33), as well as the ones identified in the present study (n=69), have originated from this ‘primordial knot’, further undergoing lineage-specific gene duplication and diversification (Figure 1 - 3). The origin and diversification of these superfamilies can be explained by a mechanism that is similar to the combinatorial peptide strategy, wherein certain venomous animals, such as cone snails, generate a remarkable diversity in their mature toxin peptides while preserving the signal and propeptide regions (18-20). Rapid events of diversification, preceded by repeated rounds of gene duplication, form the basis of the combinatorial peptide library strategy (21). These hyper-mutational events have been previously shown to be restricted to the mature peptide region of toxins (22). In contrast, the signal and propeptide regions, which are vital for the precise secretion and folding of proteins, respectively, evolve under the strong influence of negative selection pressures (23) - a molecular evolutionary trend also reported in venom coding genes of snakes (24). Spider venom coding genes appear to have followed a similar strategy. However, unlike the cone snail venom coding genes that have a recent evolutionary origin (<35 to 50 MYA; (25, 26)), spider venom toxins have likely originated from an ancestral scaffold in Opisthothelae, the MRCA of Mygalomorphae and Araneomorphae spiders, nearly 375 MYA (13). Given their significant sequence divergence since their deep-rooted evolutionary origin, the entire protein-coding gene, including the signal and propeptide regions, has accumulated significant divergence. Consistent with this hypothesis, the majority of positively selected (∼96%) identified in spider venom DRP toxins (all sites in Araneomorphae, and all but two sites in Mygalomorphae) were restricted to the mature peptide region, whereas the signal and propeptide regions harboured a minor proportion of these sites (1% and 3%, respectively; Tables S1 and S2).

### The many ways to skin a cat: innovation versus diversification of venoms

Venom is an intrinsically ecological trait that has underpinned the evolutionary success of many animals. The ability of venomous organisms to incapacitate prey and predators emanates from toxins that exhibit an array of biochemical activities and target divergent pathways. Many venomous lineages deploy a wide range of toxins from phylogenetically unrelated superfamilies. Venomous snakes, for example, have ‘recruited’ a myriad of toxins, including snake venom metalloproteinases, snake venom serine proteases, three-finger toxins, phospholipase A_2_s, L-amino acid oxidases, Kunitz-type serine protease inhibitors, kallikreins, lectins, DNases and hyaluronidases [(27, 28), Figure 5]. Similarly, spider venoms typically possess many forms of enzymes (e.g., phospholipases, proteases and chitinases), polyamines, salts and disulphide-rich toxins [(6), Figure 5]. Astonishingly, however, spider venom DRPs with diverse ion channel targeting activities, such as sodium, potassium, calcium, and chloride ion channels, predominate the venoms of nearly all spiders, constituting three-quarters of the venom (Figure 5). Phylogenetic and evolutionary assessments in this study trace the evolutionary origin of DRPs in Opisthothelae, the common ancestor of all extant spiders. This recruitment strategy, where a molecular scaffold with a single deep-rooted evolutionary origin, constitutes the major content of the venom, is unique to spiders. Venoms of most other animals are instead composed of unrelated toxin types in distinct proportions. These findings not only shed light on the fascinating evolutionary history of spider venoms but also highlight an unrealized potential of molecular scaffolds in underpinning the dramatic structural and functional diversification of the venom arsenal.

**Figure 5.**
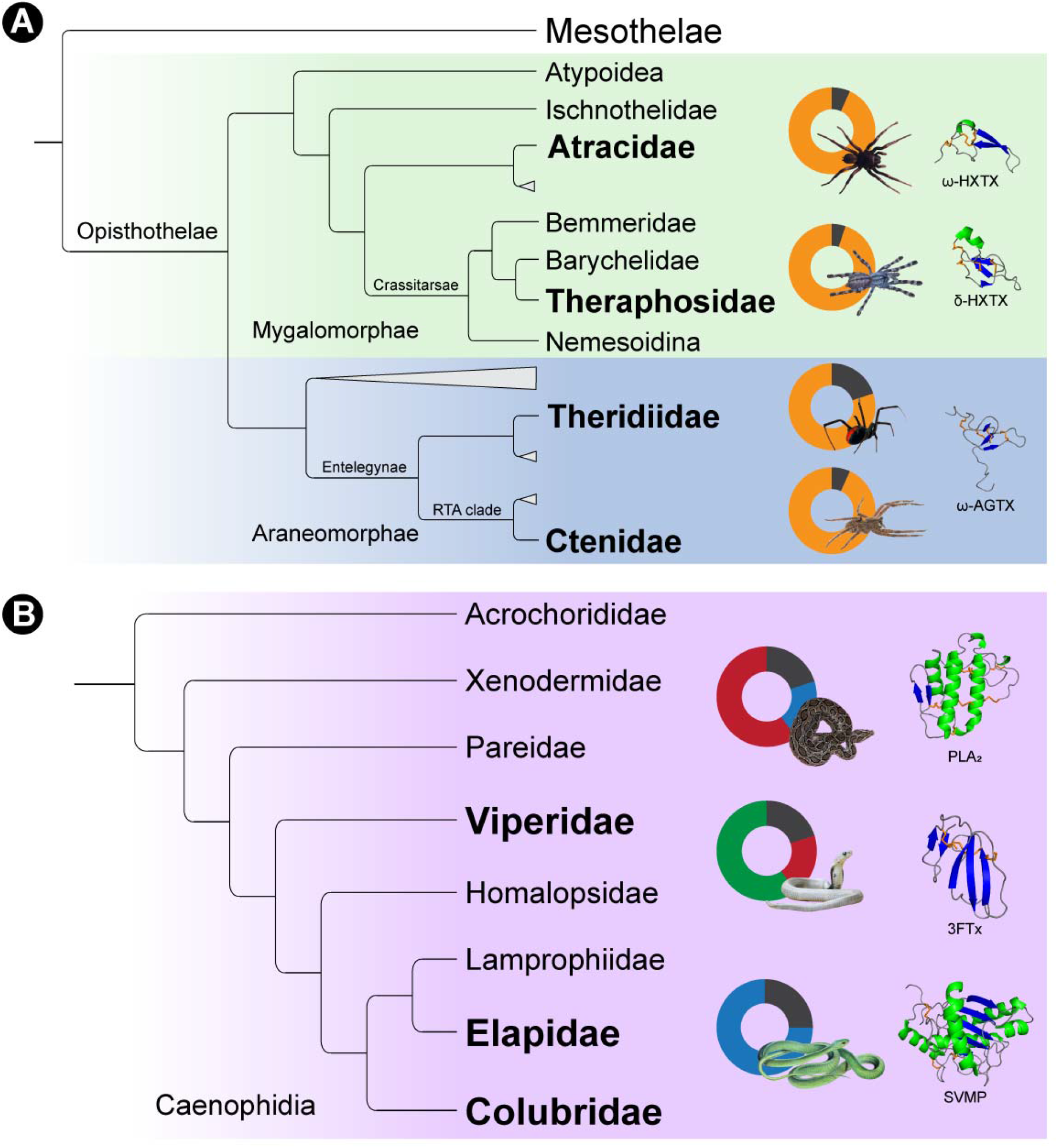
Distinct toxin scaffold recruitment strategies in spiders and snakes The figure depicts distinct toxin scaffold recruitment strategies in (A) spiders (55-57) and (B) advanced snakes (58-60). The Araneae spider phylogeny highlights the domination of disulfide-rich peptide toxins, whereas venoms of advanced snakes are constituted by diverse phylogenetically unrelated toxin superfamilies. Doughnut charts, portraying the major molecular scaffolds in venom are also shown: disulfide-rich peptides (yellow), snake venom metalloproteinases (SVMP, blue), phospholipase A_2_ (PLA_2_, red), three-finger toxins (3FTx, green) and other minor components (grey). Structures of the major scaffolds are also shown, with helices coloured in green, β-strands in blue and disulfide bonds in orange.

### Distinct prey-capture strategies have underpinned the recruitment and diversification of spider venoms

In addition to suggesting the common evolutionary origin of DRP toxins, Bayesian and maximum-likelihood phylogenies provided fascinating insights into the early diversification of DRPs in spiders. Mygalomorph DRP toxin superfamilies formed genus-specific toxin clades that suggested the recruitment of unique DRP scaffolds at the level of genera (Figure 1), while the majority of unique DRP scaffolds seemed to be recruited at the level of families in Araneomorphae (Figure 2). Only a minor fraction (6/38) of araneomorph toxin superfamilies were recruited at the genus level.

When the nature and strength of selection on venom DRPs were assessed, a strong influence of positive selection was identified on the evolution of these toxin superfamilies in mygalomorph spiders. Only a minority of these toxin superfamilies were found to be evolving under negative selection (6/19), or under near neutral evolution (1/19), while the majority (12/19) experienced diversifying selection (ω between 1.19 to 2.95; PS: 0 to 26, Figure 4). In complete contrast, the evolution of venom DRPs (13/14) in Araneomorphae was constrained by purifying selection (ω between 0.03 to 0.96; PS: 1 to 3, Figure 4), and a single superfamily was found to be evolving nearly neutrally (ω of 1.0; PS: 10). We further investigated the impact of these amino acid replacements on the structure and function of spider venom toxins. Outcomes of these evaluations suggest that the majority of replacements in mygalomorph spiders (between 0 to 29 properties) had a radical effect on the structure and/or biochemical property of the encoded toxin, while none were identified in most toxin superfamilies of Araneomorphae. Only a minor proportion of non-synonymous substitutions in two toxin superfamilies (SF40 and 68) of this lineage were reported to be radically different (Tables S1 and S2). Differences in the evolutionary histories of mygalomorph and the araneomorph DRP toxin superfamilies became apparent as we further evaluated them for the signatures of episodic diversification. We detected a greater prevalence of episodic diversifying selection on mygalomorph DRP toxin superfamilies than their araneomorph counterparts (0-34 versus 0-6 events, respectively).

Such starkly contrasting phylogenetic and evolutionary patterns are indicative of differential recruitment and diversification of DRPs in spiders. While mygalomorph spiders appear to have recruited DRPs post divergence of family members, Araneomorphae may have accomplished this before. Since most araneomorph spiders heavily rely on their foraging web for prey capture, and because these spiders mostly prey on insects (29), their venom DRPs may have become relatively less diverse (Figure 4, Table S2). In complete contrast, venom DRPs in mygalomorph spiders that mostly rely on venom, and not silk being either ambush or sit-and-wait predators, to capture a much diverse prey base, appear to have experienced a significantly greater influence of the diversifying selection [(30), Figure 4, Table S1]. These observations clearly highlight the important role of ecology and venom deployment in shaping the evolution of the spider venom arsenal. Though it should be noted that the current literature and our investigation are limited to the most diverse lineage in Araneomorphae - the RTA clade. Surprisingly, however, despite being the most speciose spider lineage, and having a significantly higher genomic diversification rate in comparison to other araneomorphs (31), the lack of toxin sequence diversity in the RTA clade is intriguing (Figure 4, Table S2). Since venom toxins from the foraging web-building araneomorphs outside the RTA clade are very poorly studied (e.g., only a handful of species are investigated from a biodiscovery perspective, and not a single toxin has been sequenced at the nucleotide level to date), the lack of venom toxin sequence diversity in the RTA clade remains intriguing and warrants further investigation.

### Deployment strategies dictate spider venom evolution

The current literature is replete with findings that support the strong influence of positive selection on genes encoding venom toxins in diverse animal lineages (32-35). Venom proteins are theorised to follow a ‘two-speed’ mode of evolution, wherein they readily diversify in animals that experience drastic shifts in ecology and/or environment - a prominent feature of evolutionarily younger lineages [e.g., cone snails and advanced snakes with evolutionary origins dating back to <35-50 MYA (36)]. This rapid expansion or the ‘expansion phase’ is shaped by a strong influence of positive selection that underpins the transition of organisms into novel ecological niches. Post these adaptive changes, the influence of diversifying selection is replaced by effects of purifying selection (the ‘purification phase’) that preserve potent toxins generated during the expansion phase. This, perhaps, explains the contrasting evolutionary regimes documented in evolutionarily younger and ancient lineages (36). Venom coding genes in evolutionarily ancient lineages are said to re-enter the expansion phase if they re-encounter dramatic shifts in ecology and environment. The only exceptions to this hypothesis are toxins that non-specifically interact with their molecular targets or those that are deployed for antipredatory defence (36). The latter hypothesis, however, mostly stems from the analyses of venom proteins that are deployed for predation. A dearth of sequence information for venom components majorly employed for antipredator defence has impeded our understanding of their evolutionary diversification.

Spiders of the genera, *Hadronyche* and *Atrax* (family Atracidae), are known to deploy their DRP toxin superfamily (SF13: Ares) predominantly for antipredatory defence (14). In contrast, tarantulas of the family Theraphosidae are known to chiefly employ their venom to capture prey animals. This provided us with a unique opportunity to comparatively investigate the molecular evolution of spider venom proteins chiefly deployed for predation (SF74, SF77, SF79, SF89, SF90, SF92 and SF99) and self-defence (SF13: Ares). Our analyses of the molecular evolutionary histories of theraphosid spider venom DRPs deployed for prey capture reveal a strong influence of diversifying selection (ω: 1.2 to 2.9; PS: 0 to 11; Table S1, Figure S11), whereas those employed for self-defence in Atracidae spiders were constrained by negative selection (ω: 0.8; PS: 3; Table S1, Figure S11). Outcomes of FUBAR and MEME analyses further corroborated these findings. FUBAR identified numerous sites (∼10%) in defensive toxins as evolving under the pervasive influence of negative selection, while MEME detected several episodically diversifying sites (∼22%) in theraphosid toxins deployed for prey capture (Table S1).

Such contrasting modes of diversification could be attributed to the ‘two-speed’ mode of venom evolution, where the offensive toxins gain an evolutionary advantage over prey by amplifying their sequence and functional diversity (36). In contrast, as defensive venoms are infrequently deployed, or have evolutionarily conserved molecular targets across predatory lineages, they experience relatively reduced effects of diversifying selection. In the absence of a need for sequence variation, purifying selection pressures instead ensure the preservation of broadly effective toxins.

## Methods

### Sequence data curation and assembly

Nucleotide datasets consisting of Mygalomorphae DRP sequences were assembled from the National Center for Biotechnology Information’s Non-redundant and Transcriptome Shotgun Assembly databases using manual search and exhaustive BLAST iterations (37). Sequences for Araneomorphae toxins were retrieved from Cole, T. J., & Brewer, M. S. (2021) (38). Translated sequences were aligned in MEGA X using MUSCLE (39, 40) before back-translation to nucleotides. Alignment was further refined by using structurally conserved cysteines as guides.

### Phylogenetic analyses

Phylogenetic histories of toxin families were reconstructed using Bayesian and maximum-likelihood inferences implemented in MrBayes 3.2.7a (41, 42) and IQ-TREE v1.6.12 (43, 44f), respectively. Bayesian analyses were run for a minimum of ten million generations using twelve Markov chains across four runs, sampling every 100th tree. Twenty-five percent of the total trees sampled were discarded as burn-in. The log-likelihood score for each tree was plotted against the number of generations to assess whether the analysis has reached an asymptote. A stop value of 0.01 was used for the average standard deviation of split frequencies. Bayesian Posterior Probability (BPP) was used to evaluate node support for the branches of Bayesian trees. ML analyses were performed using IQ-TREE with an edge-proportional partition model and 100 Bootstrap replicates. Phylogenetic trees were rooted with non-venom nucleolar cysteine-rich protein sequences from *Mastigoproctus giganteus, Stenochrus portoricensis, Prokoenenia wheeleri, Phrynus marginemaculatus* and *Cryptocellus centralis* from the class Arachnida that fall outside of the suborder Opisthothelae.

### Principal Component Analysis

PCA of signal peptide sequences from spider toxin superfamilies was performed in R [v 4.1.2; (45)] using a previously published script [(46)]. Sequences were aligned using MUSCLE in MEGA X (39, 40) and further digitising in R utilising boolean vectors. The scaled principal component values (sPC) were calculated using conventional PCA prior to plotting.

### Assessment of molecular evolution

The nature of selection shaping the evolution of DRP toxins was determined using a maximum-likelihood inference implemented in CodeML of the PAML package (47). The ratio of non-synonymous substitutions (nucleotide changes that alter the coded amino acid) to synonymous substitutions (nucleotide changes that do not alter the coded amino acid), also known as omega (ω), was estimated. A likelihood ratio test (LRT) for the nested models - M7 (null model) and M8 (alternate model) - was performed to assess the statistical significance of the findings. The Bayes Empirical Bayes (BEB) approach implemented in M8 was used to calculate the posterior probabilities for site classes (48). Amino acid sites with a posterior probability of over 95% (PP ≥ 95%) were inferred as positively selected. The episodic and pervasive nature of selection was determined using the Mixed Effect Model of Evolution (MEME) (49) and the Fast Unconstrained Bayesian AppRoximation (FUBAR) (50), respectively.

### Evaluation of selection on amino acid properties

The influence of positive selection on the biochemical and structural properties of amino acids was evaluated using TreeSAAP [v 3.2; (51)]. TreeSAAP estimates the rate of selection using a modified MM01 model (McClellan and McCracken, 2001). Statistical probabilities corresponding to a range of properties are further calculated for each amino acid. BASEML was set to run with the REV model and eight evolutionary pathway categories were defined for evolutionary pathway analyses with a sliding window size set to one. Data acquired from TreeSAAP was further visualised and processed with IMPACT_S (52).

### Structural analyses

Structural homologues of spider toxin superfamilies were identified via blast searches against the RCSB Protein Data Bank (https://www.rcsb.org/) and subsequently modelled using the SWISS-MODEL web server via user template mode (53). The resultant models were validated using MolProbity (v 4.4; https://github.com/rlabduke/MolProbity) and general Ramachandran plot. Regimes of evolutionary selection pressures were evaluated and mapped onto homology models using the Consurf webserver [(54), http://consurf.tau.ac.il/]. PyMOL v2.5.2 (Schrödinger, LLC, USA) was used to visualise and generate the images of homology models.

## Acknowledgements

This work was supported by the DBT/Wellcome Trust India Alliance Fellowship [grant number IA/I/19/2/504647] awarded to KS. NYS is thankful to Vivek Suranse and Senji Laxme R. R. (Indian Institute of Science) for insightful discussions.

## Supplementary Information for

**Fig. S1.**
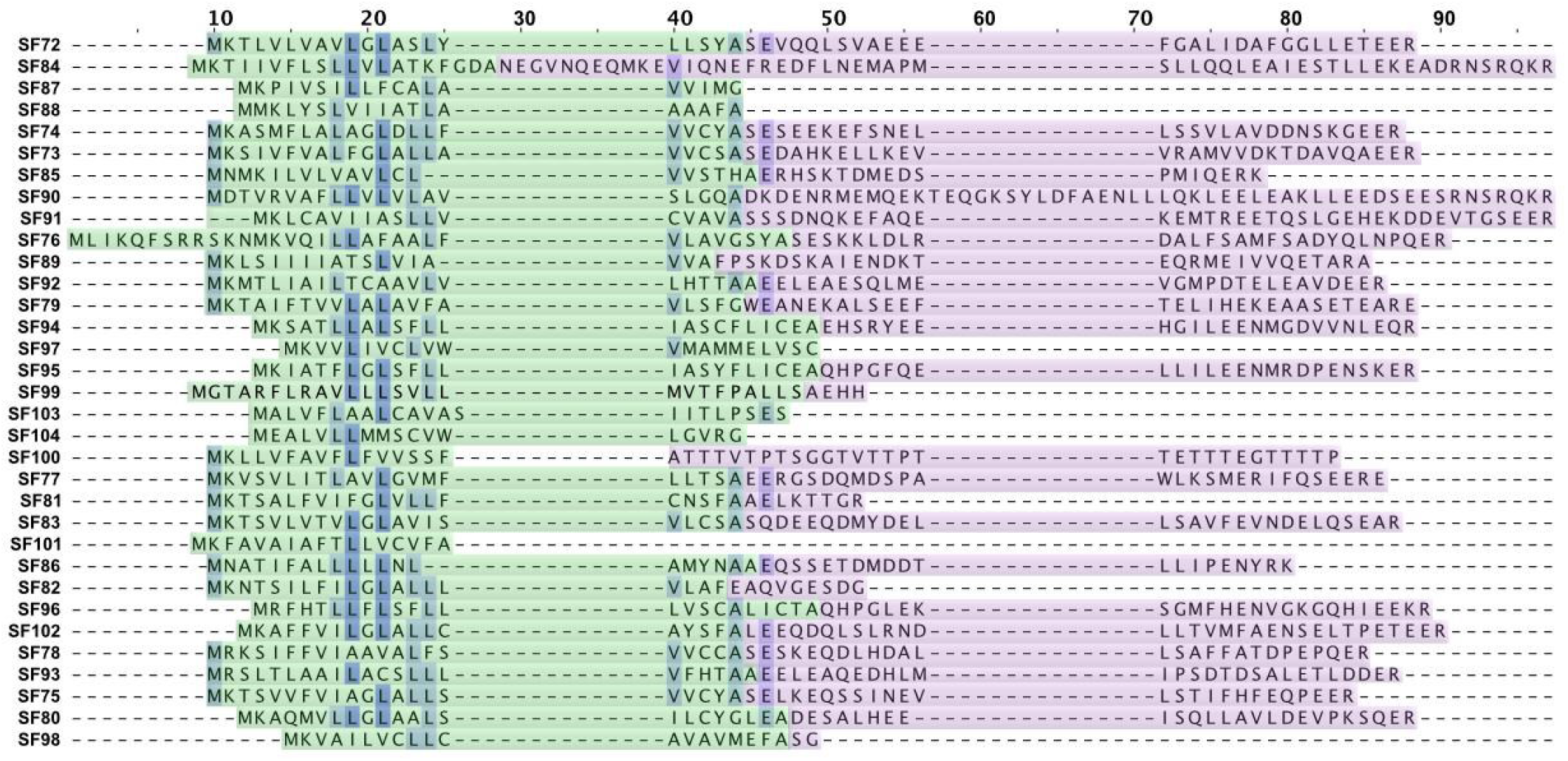
Signal peptide and propeptide alignment of novel mygalomorph superfamilies This figure shows the alignment of signal peptide and propeptide sequences from novel mygalomorph spider toxin superfamilies identified in this study. The signal peptide region is highlighted in green, while the propeptide region is represented in purple colour. Conserved sites are shaded in blue.

**Fig. S2.**
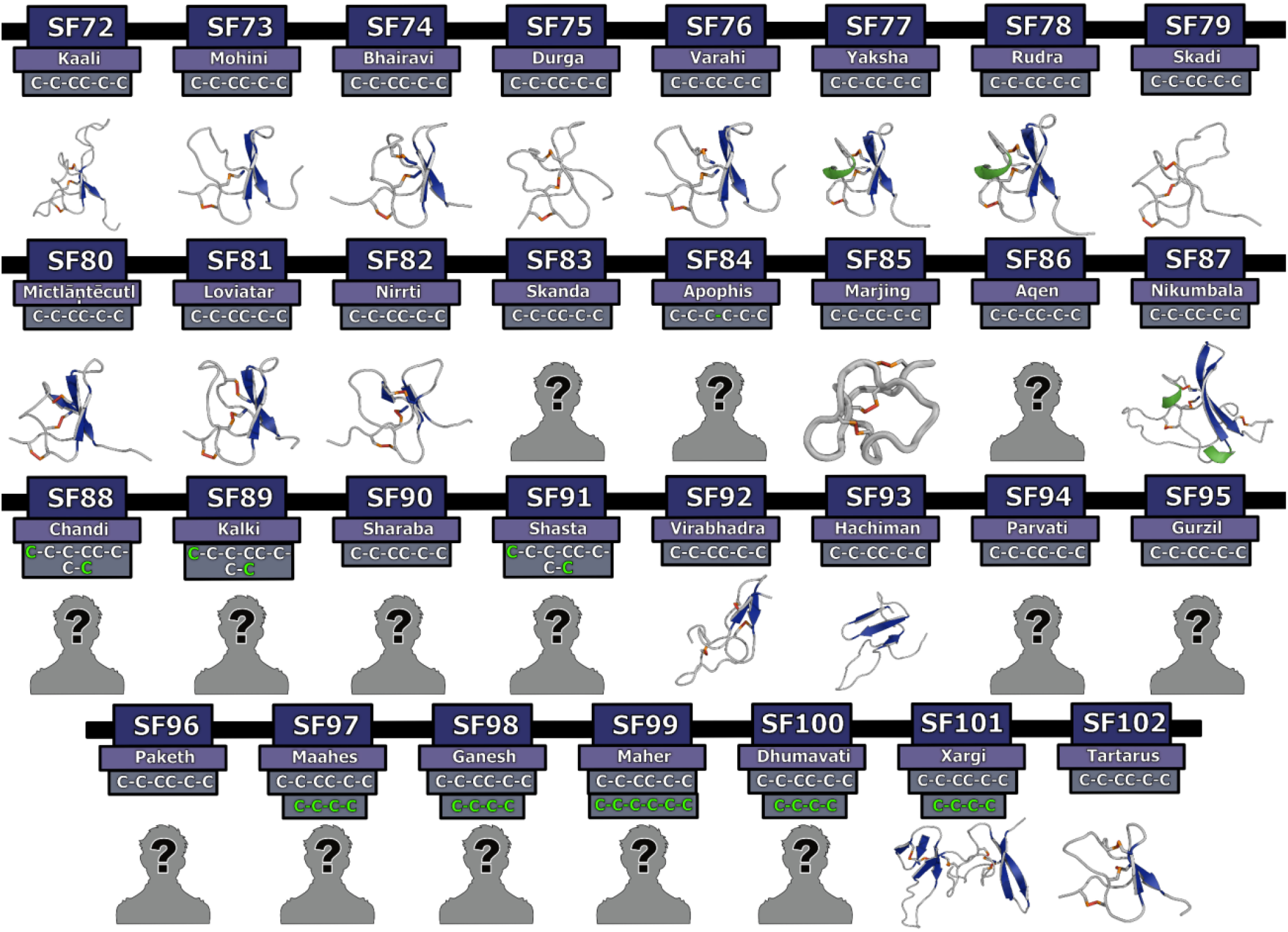
Homology models of novel Mygalomorphae toxin superfamilies This figure depicts the 3D homology models of Mygalomorphae toxin superfamilies. Here, helices are shown in green, β-strands in blue and disulfide bonds in orange. Cysteine arrangements in scaffolds are also provided above the model, where novel cysteines are shown in green text. Toxin SFs that lack structural data are indicated with a ‘?’ symbol.

**Fig. S3.**
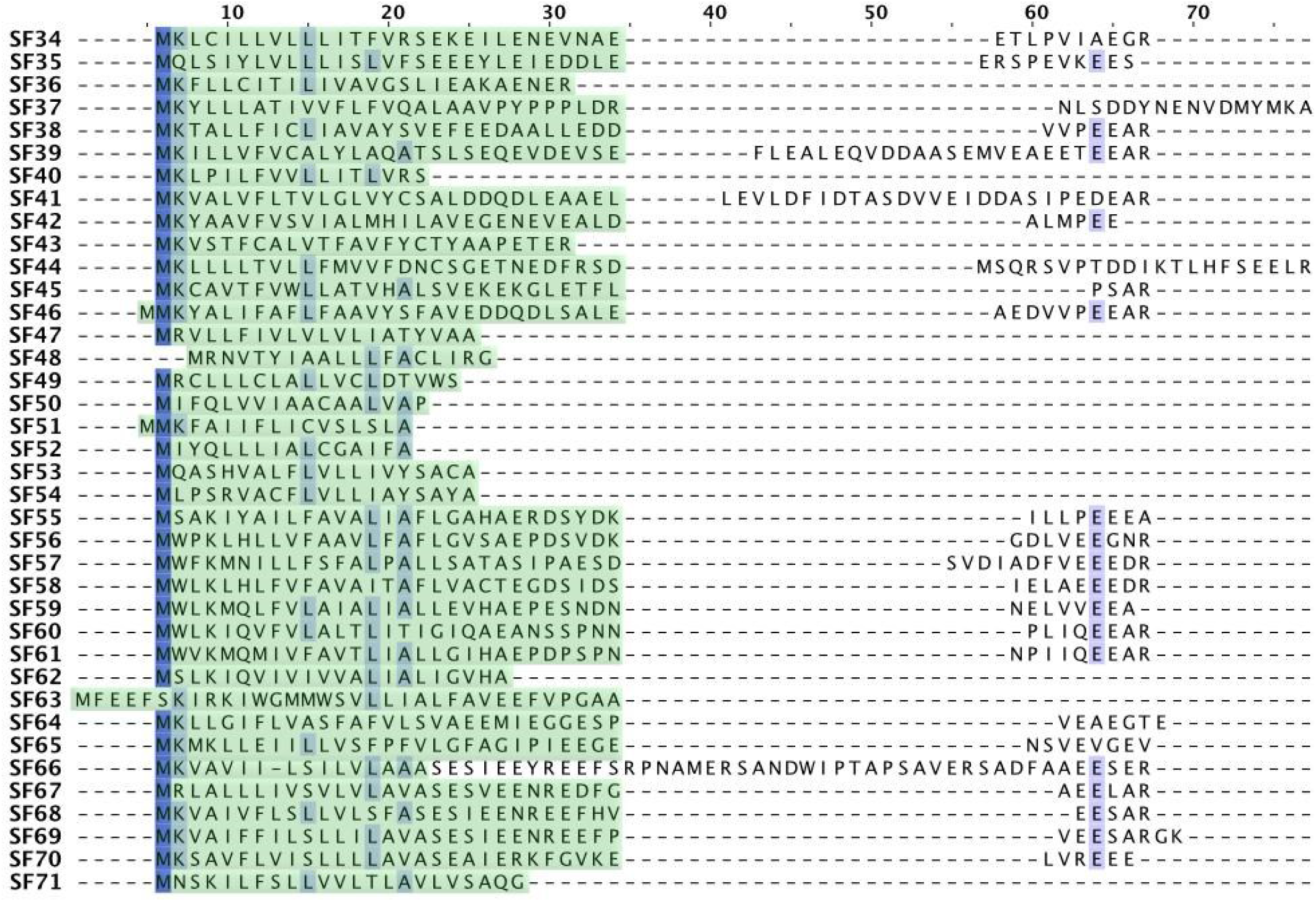
Signal peptide and propeptide alignment of araneomorph superfamilies This figure shows the alignment of signal peptide and propeptide sequences from novel araneomorph toxin superfamilies identified in this study. The signal peptide region is highlighted in green, while the conserved amino acid positions are shaded in blue. It should be noted that the propeptide region boundary could not be identified for all Araneomorphae toxin superfamilies.

**Fig. S4.**
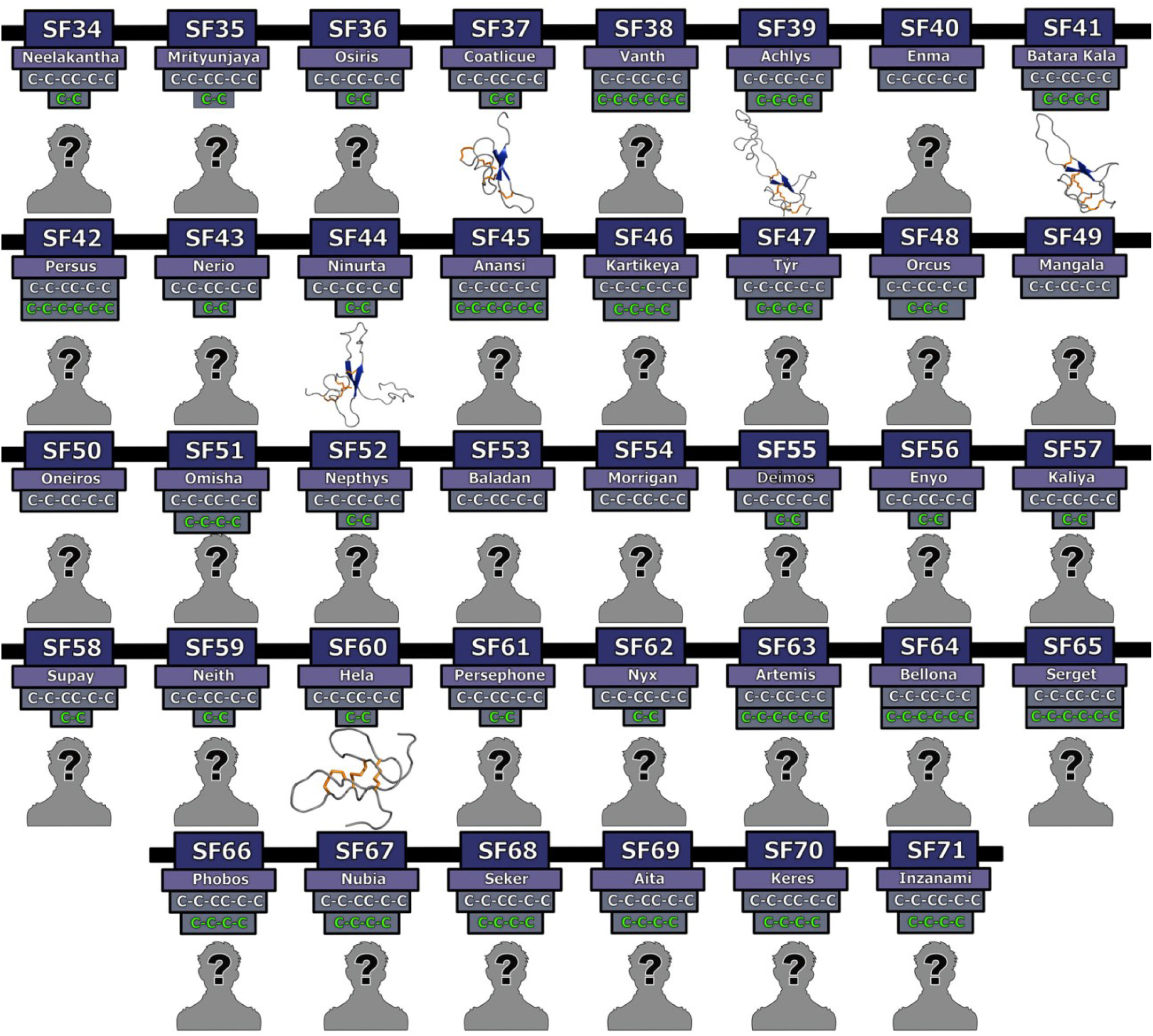
Homology models of novel Araneomorphae toxin superfamilies 3D homology models of Araneomorphae toxin superfamilies are depicted in this figure. Here, helices are shown in green, β-strands in blue and disulfide bonds in orange. Cysteine arrangements in scaffolds are also provided above the model, where novel cysteines are shown in green text. Toxin SFs that lack structural data are indicated with a ‘?’ symbol.

**Fig. S5:**
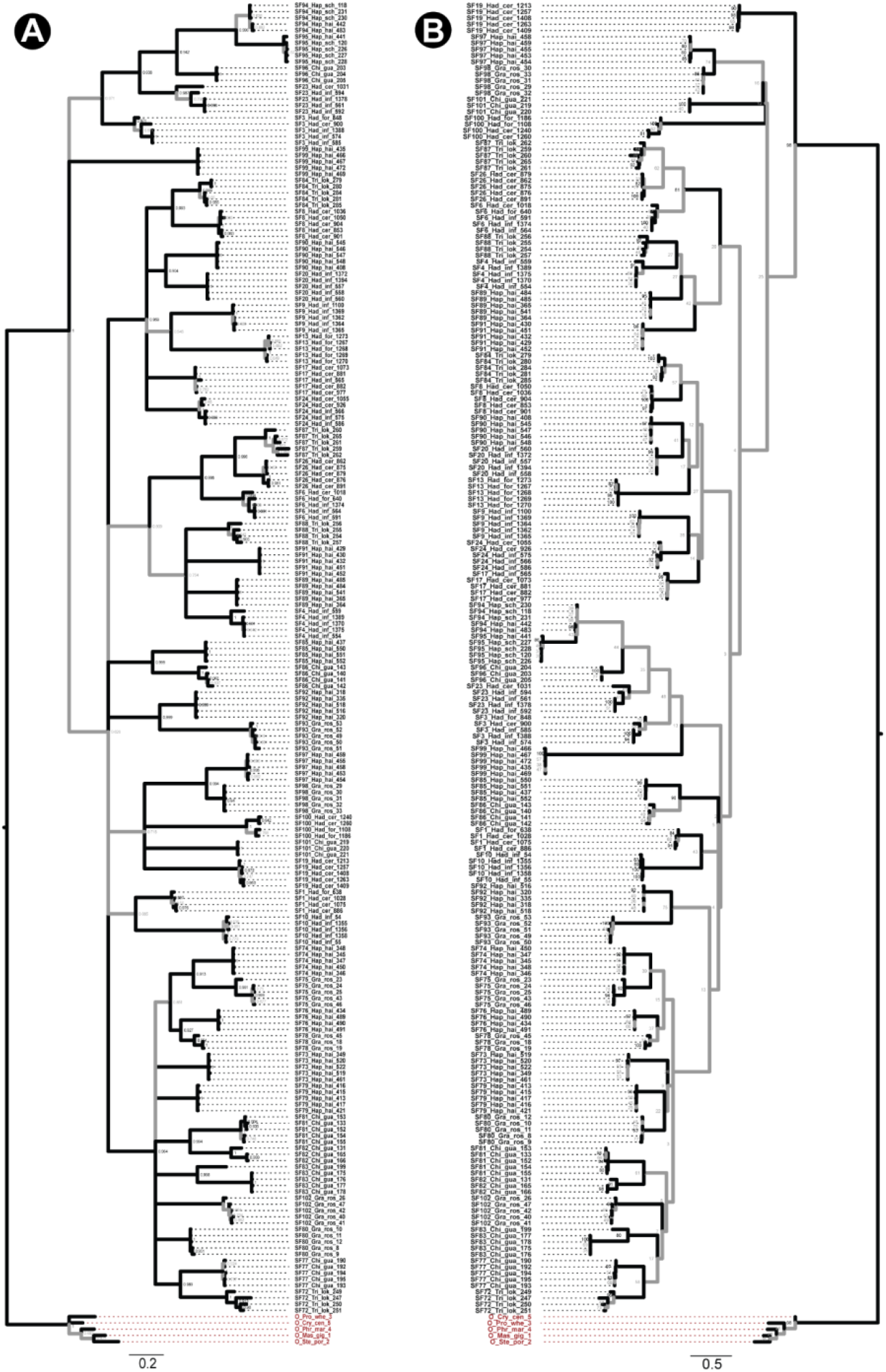
Phylogeny of Mygalomorphae spider toxin superfamilies Phylogenetic relationships of Mygalomorphae spider toxin superfamilies, assessed using Bayesian (BI; panel A) and maximum likelihood (ML; panel B) inferences, are shown in this figure. Node supports were estimated using Bayesian posterior probability (BPP) for the BI tree and bootstrapping replication (BS) for the ML tree. Branches with BPP lower than 0.95 in BI tree and BS lower than 90 in the ML tree are shown in grey. Cysteine-rich non-toxin outgroup sequences are coloured red.

**Fig. S6:**
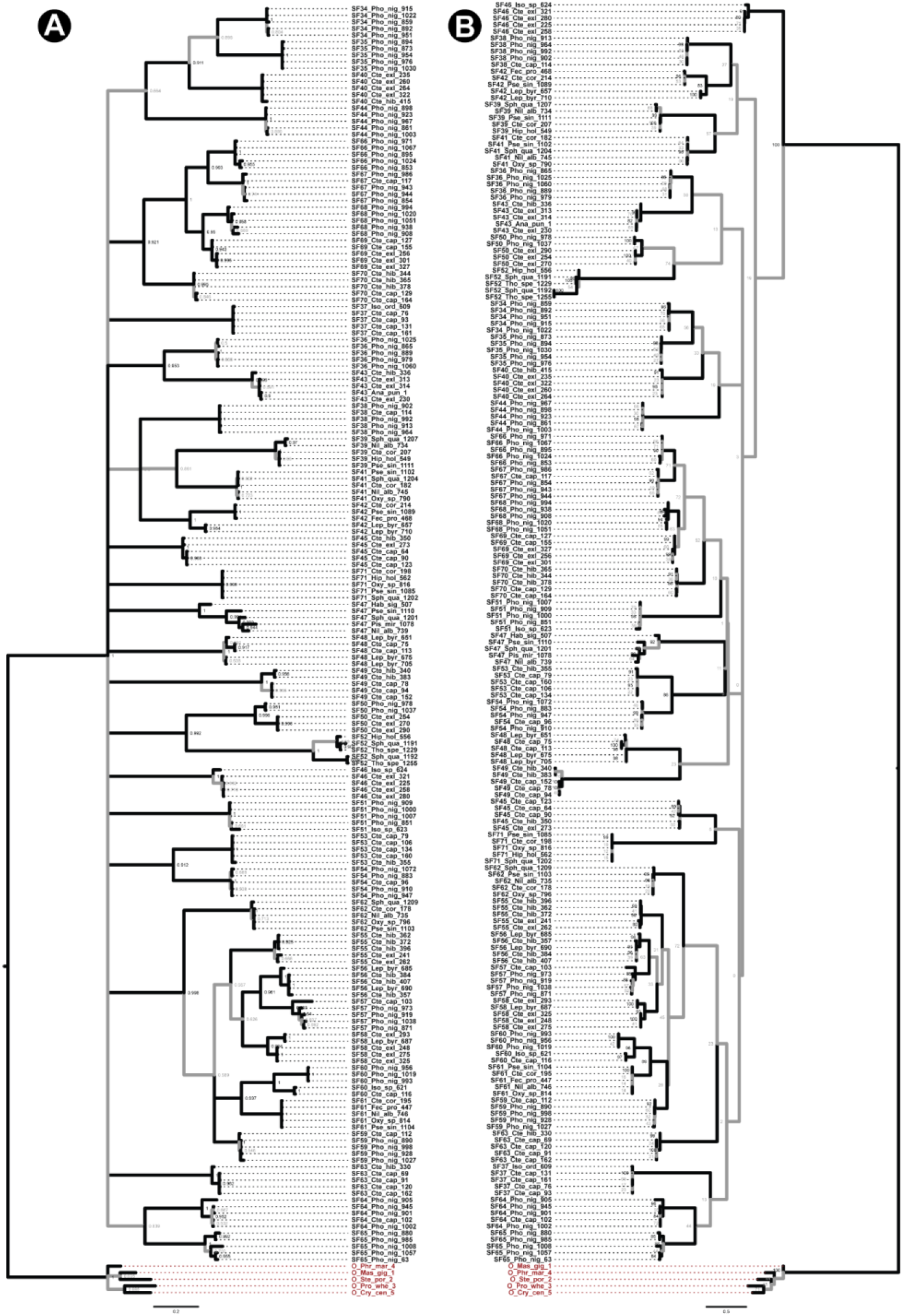
Phylogeny of Araneomorphae spider toxin superfamilies Phylogenetic relationships of Araneomorphae spider toxin superfamilies, assessed using Bayesian (BI; panel A) and maximum likelihood (ML; panel B) inferences, are shown in this figure. Node supports were estimated using Bayesian posterior probability (BPP) for the BI tree and bootstrapping replication (BS) for the ML tree. Branches with BPP lower than 0.95 in BI tree and BS lower than 90 in the ML tree are shown in grey. Cysteine-rich non-toxin outgroup sequences are coloured red.

**Fig. S7:**
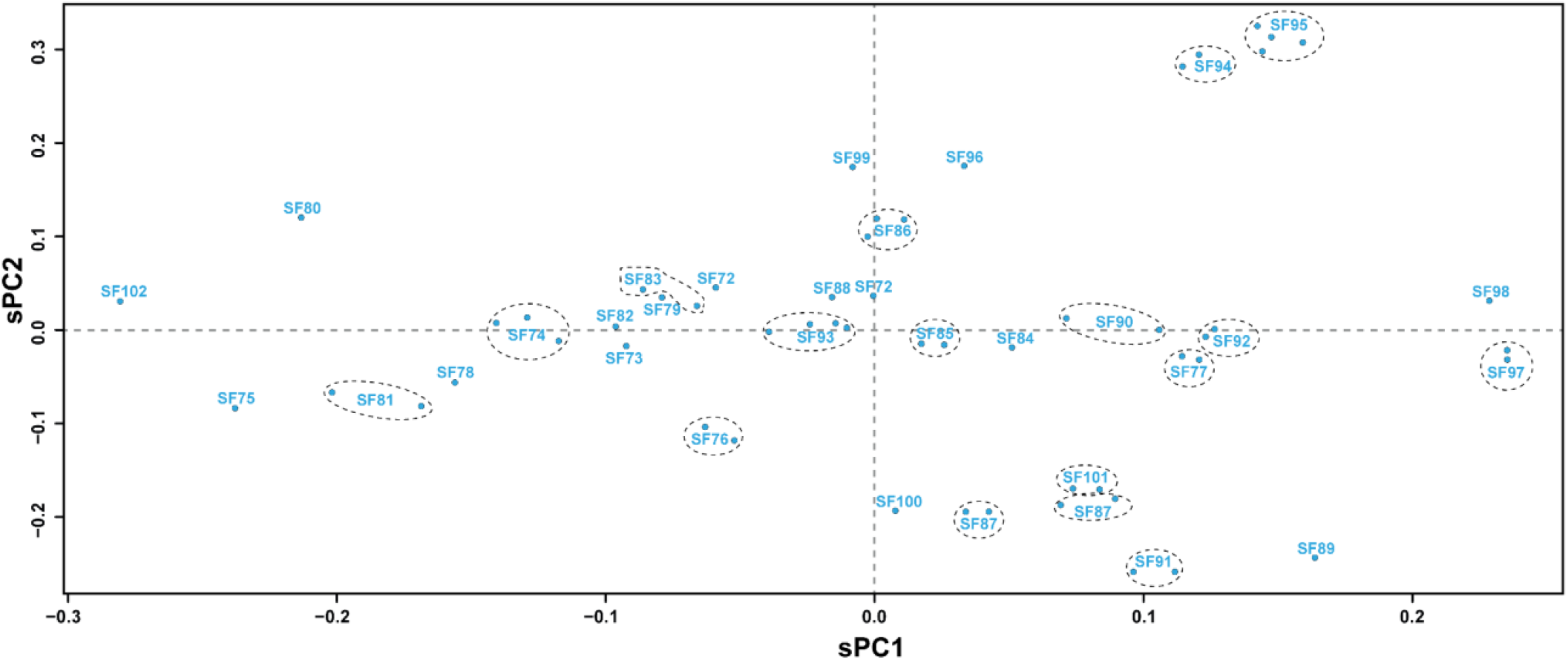
Principal component analyses for Mygalomorphae toxin superfamilies A scatter plot of scaled principal components, sPC1 and sPC2, for the signal peptide sequences of novel mygalomorph toxin superfamilies identified in this study is shown here. Signal peptide sequences belonging to a superfamily with overlapping sPC values are represented as a single dot, while others are marked with a dotted circle.

**Fig. S8:**
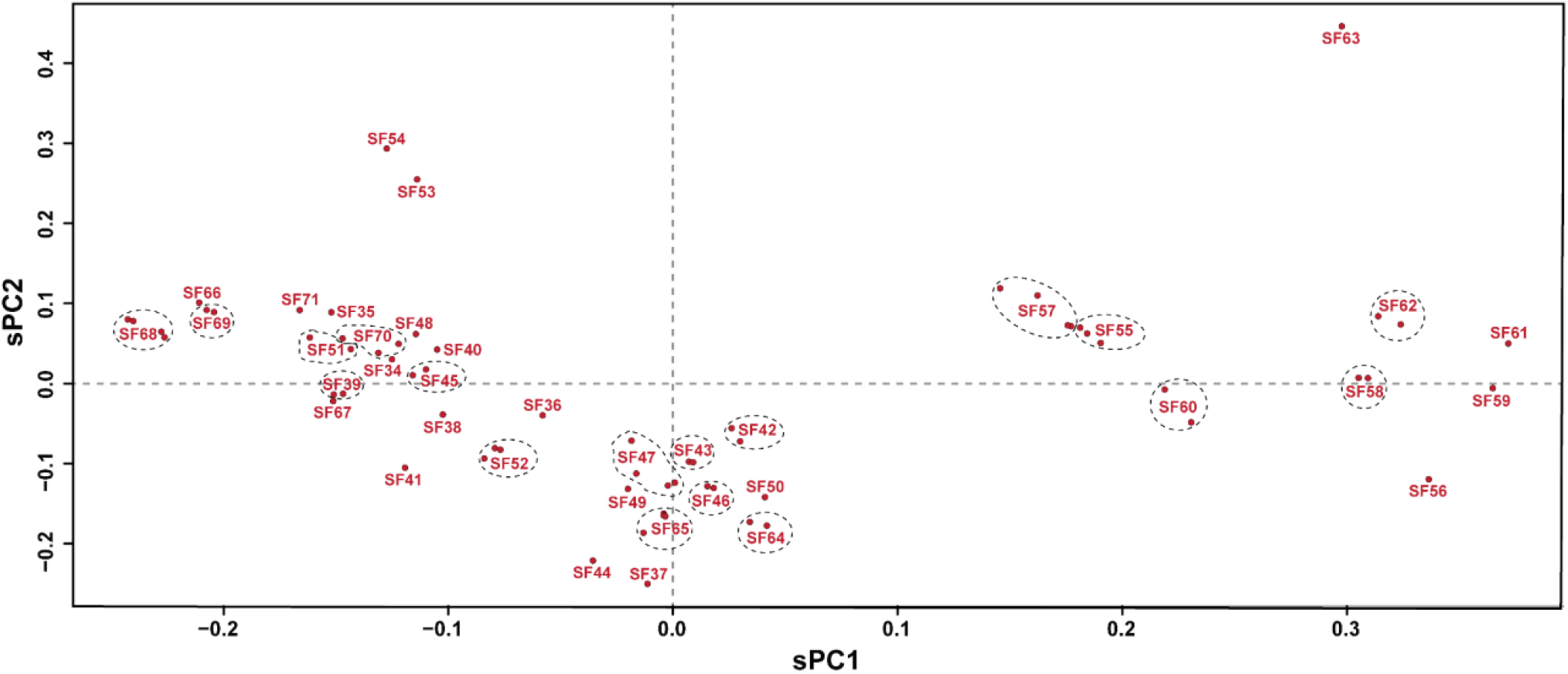
Principal component analyses for Araneomorphae toxin superfamily Scaled principal components, sPC1 and sPC2, for signal peptide sequences of novel araneomorph toxin superfamilies identified in this study are shown here in the form of a scatter plot. Signal peptide sequences belonging to a superfamily with overlapping sPC values are represented as a single dot, while others are marked with a dotted circle.

**Fig. S9:**
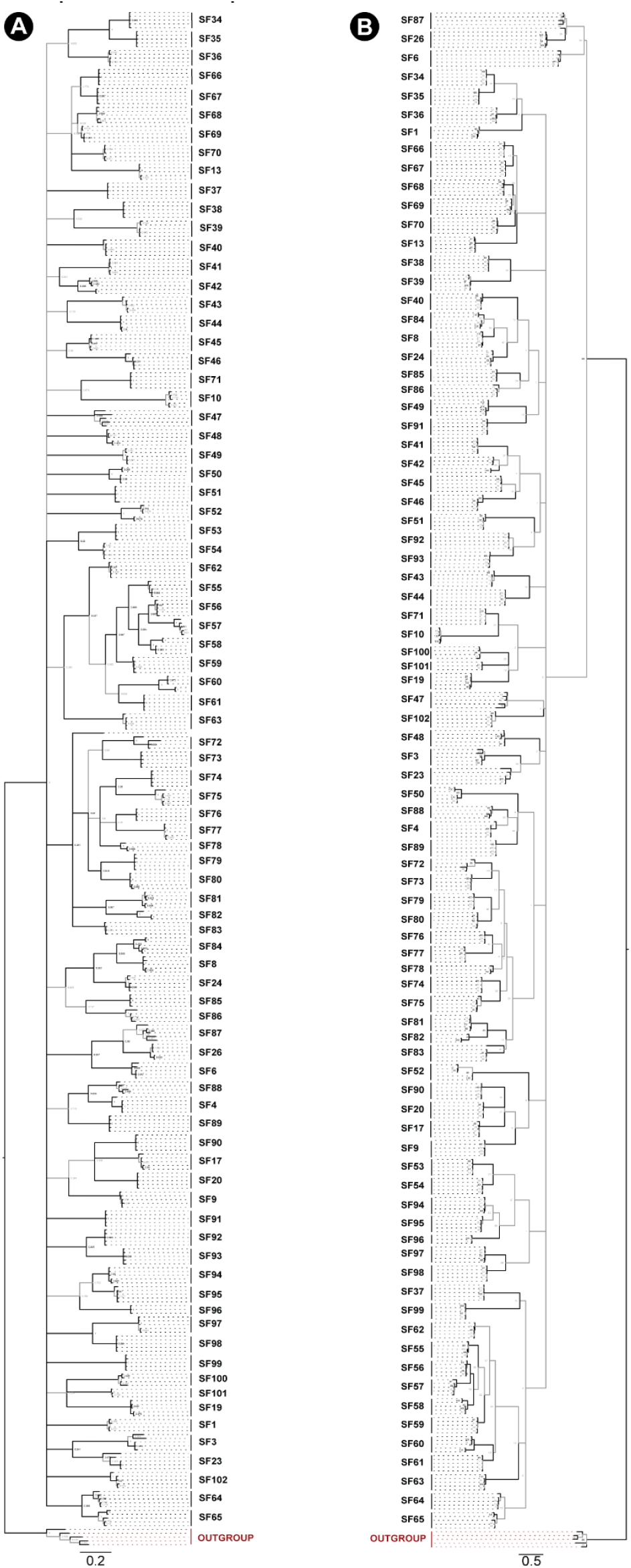
Phylogeny of Araneae spider toxin superfamilies Phylogenetic relationships of Araneae toxin superfamilies built using Bayesian (BI; panel A) and maximum likelihood (ML; panel B) inferences are shown in this figure. Node supports were estimated using Bayesian posterior probability (BPP) for the BI tree and bootstrapping replication (BS) for the ML tree. Branches with BPP lower than 0.95 in BI tree and BS lower than 90 in the ML tree are shown in grey. Cysteine-rich non-toxin outgroup sequences are coloured red.

**Fig. S10:**
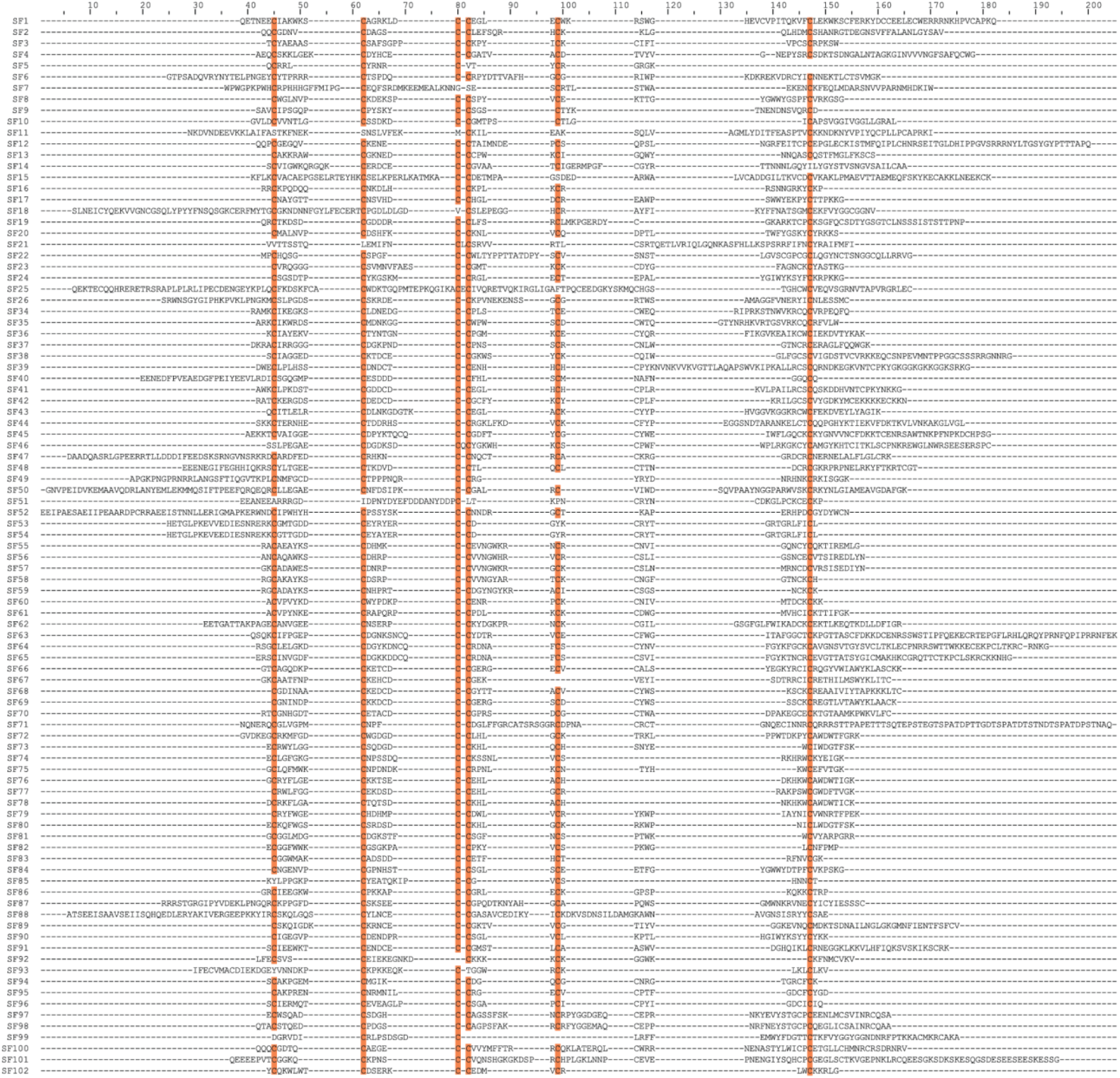
Mature peptide alignment of mygalomorph and araneomorph DRP superfamilies An alignment of mature peptide sequences from mygalomorph and araneomorph spider toxin superfamilies is shown here. Conserved amino acid positions (sequence identity ≥89%) are shaded orange.

**Fig. S11:**
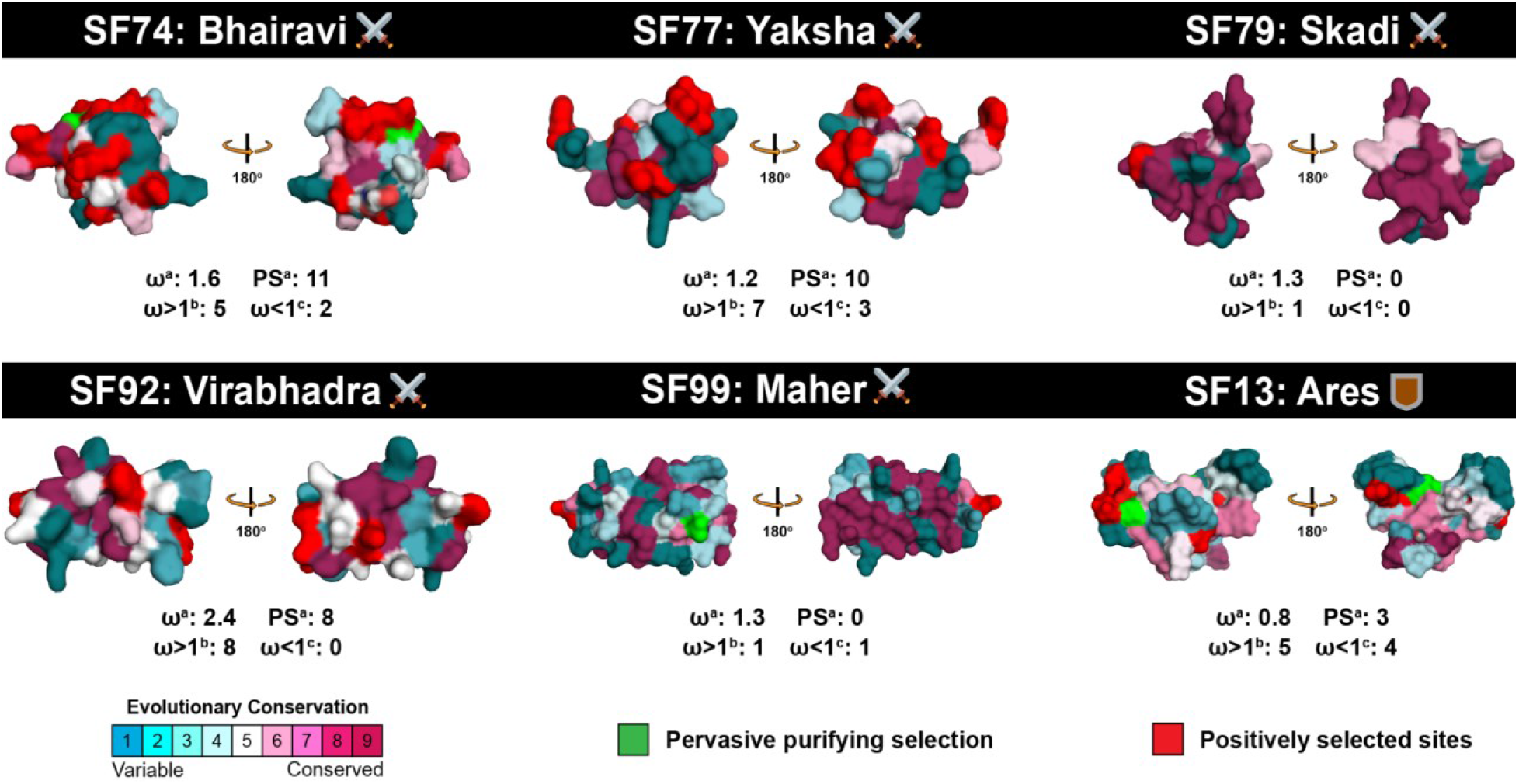
Deployment strategies dictate the evolution of spider venom This figure highlights the distinct regimes of evolutionary selection pressures acting on defensive and offensive spider venom toxin superfamilies. Positively selected sites detected by PAML (M8) and FUBAR are highlighted in red, while sites under the effect of pervasive purifying selection (FUBAR) are shown in green. A colour code indicating strength of selection is also provided. Here, ω: ratio of non-synonymous to synonymous substitutions; **a**: ω and positively selected sites (Bayes Empirical Bayes) detected by model 8 of PAML. **b**: sites experiencing pervasive influence of positive selection identified by FUBAR (ω>1); and **c**: sites experiencing pervasive influence of negative selection identified by FUBAR (ω<1).

**Table S1.** Molecular evolution of Mygalomorphae toxin superfamilies

| PAML<sup>a</sup> (M8) |  | FUBAR<sup>b</sup> | MEME<sup>c</sup> | TreeSAAP |  |
| --- | --- | --- | --- | --- | --- |
| AA | ω |  |  | Radical Property |  |
|  |  |  |  | Chemical | Physical |
| SF2 |  |  |  |  |  |
| R60 | 1.1963 | ω>1<sup>d</sup>:6<br>ω<1<sup>e</sup>:6 | 34 |  | F, Mv, Mw, V°, Ra, Hp, Ht |
| S62 |  |  |  |  | V°, Ra, Hp, an |
| L63 |  |  |  | pK | V°, Ra, Hp |
| A66 |  |  |  |  | F, Ra, Hp, an, K° |
| C68 |  |  |  |  | ac, Mv, an, K° |
| D69 |  |  |  |  | ac, K° |
| SF4 |  |  |  |  |  |
| T101 | 0.8469 | ω>1<sup>d</sup>:3<br>ω<1<sup>e</sup>:11 | 33 |  |  |
| V102 |  |  |  |  |  |
| G103 |  |  |  |  | Pα |
| G105 |  |  |  |  | Pα |
| SF7 |  |  |  |  |  |
| N89 | 2.428 | ω>1<sup>d</sup>:4<br>ω<1<sup>e</sup>:0 | 6 |  |  |
| K90 |  |  |  |  |  |
| N92 |  |  |  |  |  |
| A93 |  |  |  |  |  |
| S94 |  |  |  |  |  |
| S95 |  |  |  |  |  |
| N96 |  |  |  |  |  |
| N97 |  |  |  |  |  |
| L98 |  |  |  |  |  |
| SF9 |  |  |  |  |  |
| T73 | 0.7316 | $\omega>1^d:9$<br>$\omega<1^e:11$ | 25 | | Bl, an |
| S79 |  |  |  |  |  |
| SF10 |  |  |  |  |  |
| 0 | 0.8767 | $\omega>1^d:4$<br>$\omega<1^e:4$ | 3 | | |
| SF13 |  |  |  |  |  |
| N100 | 0.8252 | $\omega>1^d:5$<br>$\omega<1^e:4$ | 6 | | |
| G110 |  |  |  |  |  |
| L111 |  |  |  |  |  |
| SF14 |  |  |  |  |  |
| G86 | 1.9715 | $\omega>1^d:2$<br>$\omega<1^e:4$ | 34 | | |
| H90 |  |  |  |  | Ra |
| R91 |  |  |  |  |  |
| L97 |  |  |  |  |  |
| G98 |  |  |  |  |  |
| Q99 |  |  |  |  |  |
| A100 |  |  |  |  |  |
| I101 |  |  |  |  | Bl |
| L102 |  |  |  |  | Bl |
| T103 |  |  |  |  |  |
| G104 |  |  |  |  | Ra |
| F105 |  |  |  | Esm | Ra, an |
| S106 |  |  |  | h, Esm | an, RF |
| T107 |  |  |  | h | an, RF |
| V108 |  |  |  |  | an |
| T109 |  |  |  |  |  |
| N110 |  |  |  |  | RF |
| G111 |  |  |  |  | Ra, RF |
| L112 |  |  |  | h, Esm | Ra |
| S113 |  |  |  | h, Esm | Bl, an |
| A114 |  |  |  |  | Bl, an |
| I115 |  |  |  |  |  |
| F116 |  |  |  |  |  |
| C117 |  |  |  |  | RF |
| A118 |  |  |  |  | RF |
| A119 |  |  |  |  |  |
| SF17 |  |  |  |  |  |
| W100 | 1.2169 | $\omega>1^d:0$<br>$\omega<1^e:2$ | 4 | | |
| T105 |  |  |  |  | am |
| R106 |  |  |  |  | am |
| L107 |  |  |  |  |  |
| S110 |  |  |  |  |  |
| L111 |  |  |  |  |  |
| SF19 |  |  |  |  |  |
| Q19 | 1.7945 | $\omega>1^d:3$<br>$\omega<1^e:1$ | 3 | | |
| R38 |  |  |  |  |  |
| D47 |  |  |  |  |  |
| K51 |  |  |  |  |  |
| A52 |  |  |  |  |  |
| F62 |  |  |  |  |  |
| Q63 |  |  |  |  |  |
| D66 |  |  |  |  |  |
| Y68 |  |  |  |  |  |
| S70 |  |  |  |  |  |
| SF20 |  |  |  |  |  |
| S25 | 1.0695 | $\omega>1^d:0$<br>$\omega<1^e:3$ | 12 | | |
| V37 |  |  |  |  |  |
| I55 |  |  |  |  |  |
| V92 |  |  |  |  |  |
| E95 |  |  |  |  | Pc |
| P96 |  |  |  |  |  |
| T97 |  |  |  |  |  |
| L98 |  |  |  |  |  |
| T99 |  |  |  |  |  |
| W100 |  |  |  |  |  |
| Y102 |  |  |  |  |  |
| A103 |  |  |  |  |  |
| S104 |  |  |  |  |  |
| Y106 |  |  |  |  |  |
| C107 |  |  |  |  |  |
| Y108 |  |  |  |  | Pc |
| K111 |  |  |  |  |  |
| S112 |  |  |  |  |  |

| SF26 |  |  |  |  |  |
| --- | --- | --- | --- | --- | --- |
| N59 | 1.0694 | $\omega>1^d:5$<br>$\omega<1^e:13$ | 17 | | |
| E60 |  |  |  | p | Bl, Hp |
| K61 |  |  |  | p | Bl, Hp |
| E62 |  |  |  |  | Hp, an |
| S64 |  |  |  | p | Bl, Hp, an |
| S65 |  |  |  | p | Bl, Hp |
| G68 |  |  |  |  |  |
| R69 |  |  |  |  |  |
| T70 |  |  |  |  |  |
| A75 |  |  |  | p |  |
| G77 |  |  |  |  | Bl, an |
| V79 |  |  |  | p | Hp |
| N80 |  |  |  |  | an |
| E81 |  |  |  |  | an |
| SF74 |  |  |  |  |  |
| K51 | 1.6307 | $\omega>1^d:5$<br>$\omega<1^e:2$ | 8 | | |
| F54 |  |  |  |  |  |
| D55 |  |  |  |  |  |
| T58 |  |  |  |  |  |
| P33 |  |  |  |  |  |
| N67 |  |  |  |  |  |
| R68 |  |  |  |  |  |
| D72 |  |  |  |  |  |
| H74 |  |  |  |  |  |
| W79 |  |  |  |  |  |
| K80 |  |  |  |  |  |
| SF77 |  |  |  |  |  |
| M57 | 1.2384 | $\omega>1^d:7$<br>$\omega<1^e:3$ | 6 | | |
| S62 |  |  |  |  |  |
| K63 |  |  |  |  |  |
| E65 |  |  |  |  |  |
| R75 |  |  |  |  |  |
| T76 |  |  |  | Esm |  |
| F77 |  |  |  | Esm |  |
| N78 |  |  |  |  |  |
| G84 |  |  |  |  |  |
| S87 |  |  |  |  |  |
| SF79 |  |  |  |  |  |
| 0 | 1.3007 | $\omega>1^d:1$<br>$\omega<1^e:0$ | 0 | | |
| SF84 |  |  |  |  |  |
| N76 | 0.7488 | $\omega>1^d:6$<br>$\omega<1^e:7$ | 4 | | an |
| K78 |  |  |  |  |  |
| A84 |  |  |  |  | ac |
| N85 |  |  |  |  | ac |
| S87 |  |  |  |  | ac |
| P88 |  |  |  |  | ac |
| S91 |  |  |  |  |  |
| S94 |  |  |  |  |  |
| E97 |  |  |  |  | an |
| L104 |  |  |  |  |  |
| K106 |  |  |  |  |  |
| I112 |  |  |  |  |  |
| N115 |  |  |  |  |  |
| SF89 |  |  |  |  |  |
| S67 | 2.9549 | $\omega>1^d:1$<br>$\omega<1^e:0$ | 0 | | |
| S104 |  |  |  |  | Mv |
| F105 |  |  |  |  | Mv |
| S106 |  |  |  |  |  |
| C108 |  |  |  |  | an |
| V109 |  |  |  |  | an |
| SF90 |  |  |  |  |  |
| E32 | 1.5977 | $\omega>1^d:3$<br>$\omega<1^e:11$ | 1 | | |
| K38 |  |  |  |  |  |
| L60 |  |  |  |  |  |
| D63 |  |  |  |  |  |
| K112 |  |  |  |  |  |
| SF92 |  |  |  |  |  |
| V3 | 2.4954 | $\omega>1^d:8$<br>$\omega<1^e:0$ | 3 | | |
| L9 |  |  |  |  |  |
| P49 |  |  |  |  |  |
| F54 |  |  |  |  |  |
| K64 |  |  |  |  | Hp, an, K° |
| P65 |  |  |  |  | Hp, an, K° |
| K68 |  |  |  |  | ac, K° |
| V85 |  |  |  |  |  |
| SF99 |  |  |  |  |  |
| 0 | 1.3179 | $\omega > 1^d:1$<br>$\omega < 1^e:1$ | 0 | | |

**Legend:** **a**: Positively selected sites detected by the Bayes Empirical Bayes approach implemented in M8; **b**: Fast Unconstrained Bayesian AppRoximation **c**: Sites detected as experiencing episodic diversifying selection (0.05 significance) by the Mixed Effects Model Evolution (MEME). Sites detected at 0.99 and 0.95 significance are indicated in the parenthesis; **d**: number of sites under pervasive diversifying selection at the posterior probability $\geq 0.9$ (FUBAR); **e**: Number of sites under pervasive purifying selection at the posterior probability $\geq 0.9$ (FUBAR); $\omega$ : mean dN/dS.
Biochemical properties evaluated: Equilibrium Const. – ionisation, COOH (pK); Hydropathy (h); Long-range n.b. energy (EI); Polarity (p); Total n.b. energy (Et). Structural properties: $\alpha$ -helical tendencies (P $\alpha$ ); $\beta$ -structure tendencies (P $\beta$ ); Average # surrounding residues (Ns); Bulkiness (BI); Chromatographic index (RF); Coil tendencies (Pc); Compressibility (K $^\circ$ ); Helical contact energy (Ca); Mean r.m.s. fluctuation displacement. (F); Molecular volume (Mv); Molecular weight (Mw); Partial specific volume (V $^\circ$ ); Polar requirement (Pr); Power to be – C-term. $\alpha$ -helix ( $\alpha$ c); Power to be – middle, $\alpha$ -helix ( $\alpha$ m); Power to be – N-term., $\alpha$ -helix ( $\alpha$ n); Refractive index ( $\mu$ ); Solvent accessible reduct. ratio (Ra); Thermodynamics transfer hydrophobicity (Ht)

**Table S2.** Molecular evolution of Araneomorphae toxin superfamilies

| PAML<sup>a</sup> (M8) |  | FUBAR<sup>b</sup> | MEME<sup>c</sup> | TreeSAAP |  |
| --- | --- | --- | --- | --- | --- |
| AA | ω |  |  | Radical Property |  |
|  |  |  |  | Chemical | Physical |
| SF34 |  |  |  |  |  |
| L57 | 0.9494 | ω>1<sup>d</sup>:6<br>ω<1<sup>e</sup>:7 | 2 |  |  |
| R83 |  |  |  |  |  |
| SF36 |  |  |  |  |  |
| T34 | 0.9638 | ω>1<sup>d</sup>:3<br>ω<1<sup>e</sup>:6 | 2 |  |  |
| V56 |  |  |  |  |  |
| SF37 |  |  |  |  |  |
| 0 | 0.0318 | ω>1<sup>d</sup>:0<br>ω<1<sup>e</sup>:9 | 0 |  |  |
| SF38 |  |  |  |  |  |
| 0 | 0.6467 | ω>1<sup>d</sup>:0<br>ω<1<sup>e</sup>:2 | 2 |  |  |
| SF40 |  |  |  |  |  |
| D41 | 1.0142 | ω>1<sup>d</sup>:2<br>ω<1<sup>e</sup>:12 | 4 |  |  |
| I42 |  |  |  |  | αc |
| S44 |  |  |  |  |  |
| G45 |  |  |  |  |  |
| Q46 |  |  |  |  |  |
| F58 |  |  |  |  | Ns |
| N64 |  |  |  |  |  |
| N67 |  |  |  |  |  |
| G69 |  |  |  | h | αc |
| Q70 |  |  |  | h | αc |
| SF42 |  |  |  |  |  |
| 0 | 0.3161 | $\omega > 1^d:1$<br>$\omega < 1^e:12$ | 1 | | |
| SF44 |  |  |  |  |  |
| 0 | 0.8503 | $\omega > 1^d:2$<br>$\omega < 1^e:2$ | 2 | | |
| SF45 |  |  |  |  |  |
| 0 | 0.3662 | $\omega > 1^d:2$<br>$\omega < 1^e:4$ | 0 | | |
| SF46 |  |  |  |  |  |
| D103 | 0.3635 | $\omega > 1^d:3$<br>$\omega < 1^e:11$ | 0 | | |
| SF47 |  |  |  |  |  |
| 0 | 0.0861 | $\omega > 1^d:0$<br>$\omega < 1^e:57$ | 0 | | |
| SF48 |  |  |  |  |  |
| 0 | 0.1674 | $\omega > 1^d:1$<br>$\omega < 1^e:20$ | 1 | | |
| SF50 |  |  |  |  |  |
| 0 | 0.4939 | $\omega > 1^d:0$<br>$\omega < 1^e:20$ | 6 | | |
| SF51 |  |  |  |  |  |
| 0 | 0.3475 | $\omega > 1^d:2$<br>$\omega < 1^e:4$ | 0 | | |
| SF55 |  |  |  |  |  |
| 0 | 0.2892 | $\omega > 1^d:0$<br>$\omega < 1^e:2$ | 0 | | |
| SF58 |  |  |  |  |  |
| 0 | 0.5436 | $\omega > 1^d:1$<br>$\omega < 1^e:4$ | 1 | | |
| SF59 |  |  |  |  |  |
| 0 | 0.4836 | $\omega > 1^d:0$<br>$\omega < 1^e:2$ | 0 | | |
| SF60 |  |  |  |  |  |
| 0 | 0.7111 | $\omega > 1^d:2$<br>$\omega < 1^e:5$ | 2 | | |
| SF66 |  |  |  |  |  |
| G62 | 0.997 | $\omega>1^d:1$<br>$\omega<1^e:1$ | 2 | | |
| E83 |  |  |  |  |  |
| E91 |  |  |  |  |  |
| S112 |  |  |  |  |  |
| SF68 |  |  |  |  |  |
| N61 | 0.6302 | $\omega>1^d:2$<br>$\omega<1^e:6$ | 4 | | |
| I76 |  |  |  |  | ac |
| E79 |  |  |  |  |  |
| SF69 |  |  |  |  |  |
| 0 | 0.6077 | $\omega>1^d:1$<br>$\omega<1^e:10$ | 1 | | |

**Legend:** **a:** Positively selected sites detected by the Bayes Empirical Bayes approach implemented in M8 **b:** Fast Unconstrained Bayesian AppRoximation **c:** Sites detected as experiencing episodic diversifying selection (0.05 significance) by the Mixed Effects Model Evolution (MEME). Sites detected at 0.99 and 0.95 significance are indicated in the parenthesis **d:** number of sites under pervasive diversifying selection at the posterior probability $\geq 0.9$ (FUBAR) **e:** Number of sites under pervasive purifying selection at the posterior probability $\geq 0.9$ (FUBAR) $\omega$ : mean dN/dS.
Biochemical properties: Equilibrium Const. – ionisation, COOH (pK); Hydropathy (h); Long-range n.b. energy (EI); Polarity (p); Total n.b. energy (Et). Structural properties: $\alpha$ -helical tendencies (P $\alpha$ ); $\beta$ -structure tendencies (P $\beta$ ); Average # surrounding residues (Ns); Bulkiness (BI); Chromatographic index (RF); Coil tendencies (Pc); Compressibility (K $^\circ$ ); Helical contact energy (Ca); Mean r.m.s. fluctuation displacement. (F); Molecular volume (Mv); Molecular weight (Mw); Partial specific volume (V $^\circ$ ); Polar requirement (Pr); Power to be – C-term. $\alpha$ -helix (ac); Power to be – middle, $\alpha$ -helix (am); Power to be – N-term., $\alpha$ -helix (an); Refractive index ( $\mu$ ); Solvent accessible reduct. ratio (Ra); Thermodynamics transfer hydrophobicity (Ht)

**Dataset S1 (separate file)**. List of accession numbers of sequences analysed in this study with superfamily annotations.

## References

1. Lozano-Fernandez J, Carton R, Tanner AR, Puttick MN, Blaxter M, Vinther J, et al. A molecular palaeobiological exploration of arthropod terrestrialization. Philosophical Transactions of the Royal Society B: Biological Sciences. 2016;371(1699):20150133.

2. King GF. The wonderful world of spiders: preface to the special Toxicon issue on spider venoms. Elsevier; 2004. p. 471–5.

3. WSC. World Spider Catalog. Version 23.0 date accessed 08/04/2022: Natural History Museum, Bern; 2022.

4. Mullen GR, Vetter RS. Spiders (Araneae). Medical and veterinary entomology: Elsevier; 2019. p. 507–31.

5. Laxme RS, Suranse V, Sunagar K. Arthropod venoms: Biochemistry, ecology and evolution. Toxicon. 2019;158:84–103.

6. King GF, Hardy MC. Spider-venom peptides: structure, pharmacology, and potential for control of insect pests. Annual review of entomology. 2013;58:475–96.

7. Yuan C-H, He Q-Y, Peng K, Diao J-B, Jiang L-P, Tang X, et al. Discovery of a distinct superfamily of Kunitz-type toxin (KTT) from tarantulas. PloS one. 2008;3(10):e3414.

8. Wang X-h, Connor M, Smith R, Maciejewski MW, Howden ME, Nicholson GM, et al. Discovery and characterization of a family of insecticidal neurotoxins with a rare vicinal disulfide bridge. Nature structural biology. 2000;7(6):505–13.

9. Pineda SS, Chin YK-Y, Undheim EA, Senff S, Mobli M, Dauly C, et al. Structural venomics reveals evolution of a complex venom by duplication and diversification of an ancient peptide-encoding gene. Proceedings of the National Academy of Sciences. 2020;117(21):11399–408.

10. Pallaghy PK, Norton RS, Nielsen KJ, Craik DJ. A common structural motif incorporating a cystine knot and a triple-stranded β-sheet in toxic and inhibitory polypeptides. Protein Science. 1994;3(10):1833–9.

11. Undheim EA, Mobli M, King GF. Toxin structures as evolutionary tools: Using conserved 3D folds to study the evolution of rapidly evolving peptides. BioEssays. 2016;38(6):539–48.

12. Pineda SS, Sollod BL, Wilson D, Darling A, Sunagar K, Undheim EA, et al. Diversification of a single ancestral gene into a successful toxin superfamily in highly venomous Australian funnel-web spiders. BMC genomics. 2014;15(1):1–16.

13. Magalhaes IL, Azevedo GH, Michalik P, Ramírez MJ. The fossil record of spiders revisited: implications for calibrating trees and evidence for a major faunal turnover since the Mesozoic. Biological Reviews. 2020;95(1):184–217.

14. Herzig V, Sunagar K, Wilson DT, Pineda SS, Israel MR, Dutertre S, et al. Australian funnel-web spiders evolved human-lethal δ-hexatoxins for defense against vertebrate predators. Proceedings of the National Academy of Sciences. 2020;117(40):24920–8.

15. Rodríguez de la Vega RC. A note on the evolution of spider toxins containing the ICK-motif. Toxin Reviews. 2005;24(3-4):383–95.

16. Ferrat G, Darbon H. An overview of the three dimensional structure of short spider toxins. Toxin Reviews. 2005;24(3-4):359–81.

17. Chen J, Deng M, He Q, Meng E, Jiang L, Liao Z, et al. Molecular diversity and evolution of cystine knot toxins of the tarantula Chilobrachys jingzhao. Cellular and Molecular Life Sciences. 2008;65(15):2431–44.

18. Escoubas P, Rash L. Tarantulas: eight-legged pharmacists and combinatorial chemists. Toxicon. 2004;43(5):555–74.

19. Zhu S-Y, Li W-X, Zeng X-C, Liu H, Jiang D-H, Mao X. Nine novel precursors of Buthus martensii scorpion α-toxin homologues. Toxicon. 2000;38(12):1653–61.

20. Olivera BM, Hillyard DR, Marsh M, Yoshikami D. Combinatorial peptide libraries in drug design: lessons from venomous cone snails. Trends in biotechnology. 1995;13(10):422–6.

21. Sollod BL, Wilson D, Zhaxybayeva O, Gogarten JP, Drinkwater R, King GF. Were arachnids the first to use combinatorial peptide libraries? Peptides. 2005;26(1):131–9.

22. Conticello SG, Gilad Y, Avidan N, Ben-Asher E, Levy Z, Fainzilber M. Mechanisms for evolving hypervariability: the case of conopeptides. Molecular biology and evolution. 2001;18(2):120–31.

23. Duda TF, Palumbi SR. Molecular genetics of ecological diversification: duplication and rapid evolution of toxin genes of the venomous gastropod Conus. Proceedings of the National Academy of Sciences. 1999;96(12):6820–3.

24. Brust A, Sunagar K, Undheim EA, Vetter I, Yang DC, Casewell NR, et al. Differential evolution and neofunctionalization of snake venom metalloprotease domains. Molecular & Cellular Proteomics. 2013;12(3):651–63.

25. Olivera BM. EE Just Lecture, 1996: Conus venom peptides, receptor and ion channel targets, and drug design: 50 million years of neuropharmacology. Molecular biology of the cell. 1997;8(11):2101–9.

26. Duda Jr TF, Kohn AJ. Species-level phylogeography and evolutionary history of the hyperdiverse marine gastropod genus Conus. Molecular phylogenetics and evolution. 2005;34(2):257–72.

27. Casewell NR, Wüster W, Vonk FJ, Harrison RA, Fry BG. Complex cocktails: the evolutionary novelty of venoms. Trends in ecology & evolution. 2013;28(4):219–29.

28. Suranse V, Srikanthan A, Sunagar K. Animal venoms: Origin, diversity and evolution. eLS. 2018:1–20.

29. Pérez-Miles F, Perafán C. Behavior and biology of Mygalomorphae. Behaviour and ecology of spiders: Springer; 2017. p. 29–54.

30. Beydizada N, Řezáč M, Pekár S. Use of conditional prey attack strategies in two generalist ground spider species. Ethology. 2022;128(4):351–7.

31. Fernández R, Kallal RJ, Dimitrov D, Ballesteros JA, Arnedo MA, Giribet G, et al. Phylogenomics, diversification dynamics, and comparative transcriptomics across the spider tree of life. Current Biology. 2018;28(9):1489-97. e5.

32. Župunski V, Kordiš D. Strong and widespread action of site-specific positive selection in the snake venom Kunitz/BPTI protein family. Scientific reports. 2016;6(1):1–12.

33. Juarez P, Comas I, Gonzalez-Candelas F, Calvete JJ. Evolution of snake venom disintegrins by positive Darwinian selection. Molecular biology and evolution. 2008;25(11):2391–407.

34. Sunagar K, Jackson TN, Undheim EA, Ali S, Antunes A, Fry BG. Three-fingered RAVERs: Rapid Accumulation of Variations in Exposed Residues of snake venom toxins. Toxins. 2013;5(11):2172–208.

35. Sunagar K, Johnson WE, O’Brien SJ, Vasconcelos V, Antunes A. Evolution of CRISPs associated with toxicoferan-reptilian venom and mammalian reproduction. Molecular biology and evolution. 2012;29(7):1807–22.

36. Sunagar K, Moran Y. The rise and fall of an evolutionary innovation: contrasting strategies of venom evolution in ancient and young animals. PLoS genetics. 2015;11(10):e1005596.

37. Altschul SF, Gish W, Miller W, Myers EW, Lipman DJ. Basic local alignment search tool. Journal of molecular biology. 1990;215(3):403–10.

38. Cole TJ, Brewer MS. Killer Knots: Molecular evolution of Inhibitor Cystine Knot toxins in wandering spiders (Araneae: Ctenidae). bioRxiv. 2021.

39. Edgar RC, Batzoglou S. Multiple sequence alignment. Current opinion in structural biology. 2006;16(3):368–73.

40. Kumar S, Stecher G, Li M, Knyaz C, Tamura K. MEGA X: molecular evolutionary genetics analysis across computing platforms. Molecular biology and evolution. 2018;35(6):1547.

41. Altekar G, Dwarkadas S, Huelsenbeck JP, Ronquist F. Parallel metropolis coupled Markov chain Monte Carlo for Bayesian phylogenetic inference. Bioinformatics. 2004;20(3):407–15.

42. Ronquist F, Teslenko M, Van Der Mark P, Ayres DL, Darling A, Höhna S, et al. MrBayes 3.2: efficient Bayesian phylogenetic inference and model choice across a large model space. Systematic biology. 2012;61(3):539–42.

43. Nguyen L-T, Schmidt HA, Von Haeseler A, Minh BQ. IQ-TREE: a fast and effective stochastic algorithm for estimating maximum-likelihood phylogenies. Molecular biology and evolution. 2015;32(1):268–74.

44. Chernomor O, Von Haeseler A, Minh BQ. Terrace aware data structure for phylogenomic inference from supermatrices. Systematic biology. 2016;65(6):997–1008.

45. R Development Core Team. R: A language and environment for statistical computing. 4.2.0 ed: R Foundation for Statistical Computing; 2021.

46. Konishi T, Matsukuma S, Fuji H, Nakamura D, Satou N, Okano K. Principal component analysis applied directly to sequence matrix. Scientific Reports. 2019;9(1):1–13.

47. Yang Z. PAML 4: phylogenetic analysis by maximum likelihood. Molecular biology and evolution. 2007;24(8):1586–91.

48. Yang Z, Wong WS, Nielsen R. Bayes empirical Bayes inference of amino acid sites under positive selection. Molecular biology and evolution. 2005;22(4):1107–18.

49. Murrell B, Wertheim JO, Moola S, Weighill T, Scheffler K, Kosakovsky Pond SL. Detecting individual sites subject to episodic diversifying selection. PLoS genetics. 2012;8(7):e1002764.

50. Murrell B, Moola S, Mabona A, Weighill T, Sheward D, Kosakovsky Pond SL, et al. FUBAR: a fast, unconstrained bayesian approximation for inferring selection. Molecular biology and evolution. 2013;30(5):1196–205.

51. Woolley S, Johnson J, Smith MJ, Crandall KA, McClellan DA. TreeSAAP: selection on amino acid properties using phylogenetic trees. Bioinformatics. 2003;19(5):671–2.

52. Maldonado E, Sunagar K, Almeida D, Vasconcelos V, Antunes A. IMPACT_S: integrated multiprogram platform to analyze and combine tests of selection. PloS one. 2014;9(10):e96243.

53. Waterhouse A, Bertoni M, Bienert S, Studer G, Tauriello G, Gumienny R, et al. SWISS-MODEL: homology modelling of protein structures and complexes. Nucleic acids research. 2018;46(W1):W296–W303.

54. Ashkenazy H, Abadi S, Martz E, Chay O, Mayrose I, Pupko T, et al. ConSurf 2016: an improved methodology to estimate and visualize evolutionary conservation in macromolecules. Nucleic acids research. 2016;44(W1):W344–W50.

55. Palagi A, Koh JM, Leblanc M, Wilson D, Dutertre S, King GF, et al. Unravelling the complex venom landscapes of lethal Australian funnel-web spiders (Hexathelidae: Atracinae) using LC-MALDI-TOF mass spectrometry. Journal of proteomics. 2013;80:292–310.

56. Oldrati V, Koua D, Allard P-M, Hulo N, Arrell M, Nentwig W, et al. Peptidomic and transcriptomic profiling of four distinct spider venoms. PloS one. 2017;12(3):e0172966.

57. Diniz MR, Paiva AL, Guerra-Duarte C, Nishiyama Jr MY, Mudadu MA, Oliveira Ud, et al. An overview of Phoneutria nigriventer spider venom using combined transcriptomic and proteomic approaches. PloS one. 2018;13(8):e0200628.

58. Faisal T, Tan KY, Tan NH, Sim SM, Gnanathasan CA, Tan CH. Proteomics, toxicity and antivenom neutralization of Sri Lankan and Indian Russell’s viper (Daboia russelii) venoms. Journal of Venomous Animals and Toxins including Tropical Diseases. 2021;27.

59. Dutta S, Chanda A, Kalita B, Islam T, Patra A, Mukherjee AK. Proteomic analysis to unravel the complex venom proteome of eastern India Naja naja: Correlation of venom composition with its biochemical and pharmacological properties. Journal of proteomics. 2017;156:29–39.

60. Pla D, Sanz L, Whiteley G, Wagstaff SC, Harrison RA, Casewell NR, et al. What killed Karl Patterson Schmidt? Combined venom gland transcriptomic, venomic and antivenomic analysis of the South African green tree snake (the boomslang), Dispholidus typus. Biochimica et Biophysica Acta (BBA)-General Subjects. 2017;1861(4):814–23.

